# PLANT WATER-CARBON RELATIONS IN HUANGLONGBING-AFFECTED CITRUS TREES AND THE IMPACT OF HLB EXPOSURE DURATION, FRUIT DEVELOPMENTAL STAGE, AND SOURCE-SINK MANIPULATION ON THESE RELATIONS

**DOI:** 10.64898/2026.10.01.755948

**Authors:** Dorcas O. Alade, Tripti Vashisth, Christopher I. Vincent

## Abstract

Bulk phloem flow is driven by pressure gradients generated by water drawn from the xylem, coupling source-sink transport with water status. Huanglongbing (HLB), a phloem-affecting bacterial disease of citrus, disrupts phloem transport, and causes canopy water deficit. Although improved water management can enhance HLB-affected tree health, the effects of HLB on water-driven source-sink relations are unknown. Fruits function as both carbon sink and water reservoir, but the role of fruit in regulating whole-tree water status remain poorly understood. We examined leaf-fruit water relations in HLB-affected citrus and evaluated how HLB exposure duration, fruit developmental stage, and source-sink manipulation shape these dynamics. Source-sink relations were manipulated in branches of symptomatic trees with long-term (four years) and recent (six months) HLB exposure. We girdled branches with a single fruit as the sink and defoliated at (0%, 50%, or 75%), and an ungirdled, non-defoliated branch served as control. We measured stomatal conductance, leaf hydraulic traits, leaf and fruit soluble sugars, diurnal fruit shrinkage (to estimate xylem backflow) and fruit water loss. Source-sink manipulation did not affect shoot total soluble sugar. HLB-exposure resulted in less negative osmotic potential, more negative turgor potential and reduced modulus of elasticity and relative water content at turgor loss point. HLB exposure also increased diurnal fruit shrinkage, with shrinkage responses to source-sink manipulation varying across fruit development stages. Collectively, these findings suggest that long-term HLB exposure promotes leaf acclimation to water deficits, however, high sink demand may diminish the leaf water status regulation at Stage II, when fruit sink carbon demand is highest.

## INTRODUCTION

Plant water and carbon dynamics are tightly coupled. Water availability regulates carbon gain by affecting stomatal conductance and thus CO₂ uptake, and carbon allocation and transport depend on adequate hydraulic supply to sustain phloem pressure gradients and metabolic demand (Xiong and Nadal 2020; Huang *et al*., 2024). Plants acquire water primarily through root uptake from the soil, but the loss, movement, retention, and redistribution of that water within the plant are influenced by stomatal regulation and osmotic adjustment, as well as redistribution from stems, and even fruits through xylem backflow contributing to leaf water supply (Goldschmidt and Koch, 1996; Cardoso *et al*., 2020).

Xylem backflow exemplifies reciprocal exchange of water and carbon between fruits and leaves. When leaf transpiration exceeds water supply coming from the roots depending on plant water status, fruit lose water to the transpiring leaves (Yasunaga *et al*., 2013). Xylem backflow from fruit has been reported to supply the water that generates pressure for phloem carbon transport to sinks including fruit (Goldschmidt and Koch, 1996). The supply of water directly influences carbon dynamics by determining leaf water potential, turgor maintenance, and the efficiency of phloem loading, transport, and unloading (Stanfield and Bartlett, 2022). Conversely, carbon metabolism feeds back on plant water relations through osmotic adjustment, changes in cell wall elasticity, and sink-driven modulation of transpiration and hydraulic conductance. Thus, disruptions to either water or carbon pathways can therefore alter source-sink balance, reduce transport efficiency, and ultimately constrain whole-plant performance (Hamido *et al*., 2017; Kim *et al*. 2021; Chen *et al*., 2025).

In citrus, an evergreen tropical tree, carbon transport is disrupted by huanglongbing (HLB), a phloem-limited disease. HLB is a global threat to citrus industries and is caused by *Candidatus* ‘Liberibacter asiaticus’ (*C*Las), which is vectored by the Asian citrus psyllid, (*Diaphorina citri* [Kuwayama]) (Halbert & Manjunath, 2004; Welker *et al*., 2022). Disruption of phloem-mediated carbon transport under HLB has been associated with declines in root activity, including reductions in feeder root density and shortened root lifespan, ultimately leading to root dieback (Johnson *et al*., 2014; Sutton and Vashisth, 2025). This loss of functional roots limits water uptake and predisposes HLB-affected trees to water deficit, (Khalilzadeh *et al*., 2026), which may further exacerbate carbon transport, given the strong dependence of phloem transport on plant water status.

Leaf water potential serves as a central indicator of plant water status and determines the direction of water movement within the plant (Jarvis *et al*., 1976; Ali *et al*., 2023). Leaf water potential is determined by water availability and xylem resistance (Turner, 1981). It is strongly influenced by leaf solute concentration, indicating sucrose which generates negative osmotic potential. This osmotic adjustment promotes water influx into leaf cells, increasing turgor pressure and sustaining the pressure gradients required for continued sugar transport (Ali *et al*., 2023). Thus, leaf water potential, sucrose concentration, and turgor pressure function as an interdependent system that coordinates carbon export with water balance.

However, under water deficit, trees may lose the capacity to maintain adequate turgor pressure. In HLB-affected trees, where source-sink imbalance and water deficit have been reported (Hamido *et al*., 2017; Chen *et al*., 2025), sucrose accumulates in leaves without a corresponding decline in leaf water potential (da Silva *et al*., 2021). Though the relative impact on turgor potential is unknown, this decoupling suggests that osmotic regulation is impaired and that leaves may rely on alternative water sources, potentially including fruit xylem backflow, to sustain leaf hydration and hydraulic function.

Despite these impairments, reports indicate that prolonged exposure to HLB can be accompanied by partial physiological acclimation of carbon dynamics, including recovery of leaf gas exchange (net photosynthesis, stomatal conductance) relative to trees with recent HLB infection (Keeley *et al*., 2022). In addition, HLB-tolerant cultivars have been documented to increase xylem vessel area which is expected to increase water conductivity (Robledo *et al*., 2024), and improved water-management practices such as shading improved leaf water status, greater photosynthesis and lesser starch accumulation which suggests improved carbon transport (Suh *et al*., 2021; Khalilzadeh et al., 2025). Sutton and Vashisth (2025) also reported that frequent low-volume irrigation enhances tree health of HLB-affected trees. These improvements may reflect enhanced turgor driven by better water availability, which could partially sustain carbon transport despite phloem impairment. Improved water management and long-term exposure might suggest acclimation to HLB and water deficit, however the physiological mechanisms underlying these responses, particularly how leaf osmotic adjustment, turgor pressure and compensation from xylem backflow interact remain poorly understood.

Therefore, the objective of this study was to examine leaf-fruit water relations in HLB-affected citrus and determine how HLB exposure duration, and source-sink manipulation shape these dynamics. Because these dynamics may be affected by fruit development, we repeated the work through the three major fruit development stages. We hypothesized that: (1) HLB increases solute concentration which decreases solute potential and thus increases leaf turgor potential and; (2) HLB increases reliance on fruit xylem backflow as a compensatory mechanism, which is further exacerbated by defoliation. Our findings provide quantitative knowledge of the physiological coupling between water and carbon dynamics in trees affected by phloem disease.

## MATERIALS AND METHODS

### Study site and Plant material selection

The experiment was conducted using field grown, 4-years-old ‘Valencia’ sweet orange [*Citrus* × *sinensis* (L.) Osbeck] scion grafted onto ‘Kuharske’ citrange [*Citrus sinensis* (L.) Osbeck × *Poncirus trifoliata* (L.) Raf.] rootstock, located in University of Florida Citrus Research and Education Center, Lake Alfred, Florida, USA with latitude and longitude coordinates of 28°06′22″ N, 81°42′54″ W in an Entisol soil that is about 96% Candler fine sand. Trees were under micro-sprinkler irrigation, and the orchard was maintained according to University of Florida Institute of Food and Agricultural Sciences recommendations. The weather pattern differed at different stages of fruit development; maximum temperatures were higher for Stages I and II (May and July 2024; about 36 °C) and lower for Stage III (November; 30°C) and the rainfall of 6.84, 12.80 and 1.04 cm for Stages I, II and III respectively (Supplementary Table S1). The experimental plots were part of an experiment addressing planting establishment methods, previously described in Chuang *et al*. (2024). The experiment resulted in plants that were exposed to HLB infection at different times, but which by the time of the present work were growing under the same conditions, which we took advantage of for this study.

For the long-exposed treatment, we used plots that had been established using a metalized UV- reflective mulch, which can reduce infestation by insects like psyllids, increases water/nutrient use efficiency and light distribution within the canopy (Ghoveisi *et al*., 2023). The reflective mulch had broken down after two years, approximately two years prior to this study. However, of the treatments in the study in which plants became infected (those established using individual protective covers), this group showed the strongest canopy growth, which is why we chose these trees for the present work.

For the recently-exposed treatment, we used trees that had been established using individual protective covers (IPCs). IPCs are polyethylene mesh bags with small pore diameter to prevent psyllid infestation of citrus trees, thereby protecting them from infection with ‘*Ca*. L. asiaticus’. IPCs have been reported as effective protection against psyllids while enhancing tree growth (Gaire *et al*., 2024), until the trees have grown to fill the inside of the enclosure, at which point the IPCs must be removed, exposing the tree to infection with the now-endemic disease (Albrecht *et al*., 2025). The trees were established four years before the start of this study with the reflective mulch plots having been exposed for HLB due to psyllid feeding since inception. The IPCs were removed from the trees six months before the start of this study exposing them more recently to HLB infection. Based on previous studies indicating that HLB infection is typically established within six months of exposure, trees exposed for at least six months were are presumed to be infected because of high vector pressure (Pierre et al., 2021). Additionally, HLB symptoms were observed on all trees. Thus, in this experiment trees from the IPC plots are termed recently-exposed trees while trees from the reflective mulch plots are termed long- exposed trees. Ten single-tree replicates of each category were randomly selected for the purposes of this study.

### Source-sink manipulation treatment

To assess the responses of the two disease categories to source-sink manipulation, on each tree, four different branches with one fruit each were tagged to achieve four different defoliation ratios as described in Fig.1 and as follows. One branch was kept intact as a control treatment (no girdling or defoliation), three were girdled to separate them from alternate sinks or sources, and the girdle was wrapped with parafilm to prevent it from desiccating. One of the girdled branches was left intact without defoliation (0% defoliation) and two of the girdled branches were defoliated to manipulate the source-sink ratio at 50% or 75% of the original leaf number. Measurements were performed between seven to fifteen days (depending on the measurement) after the source-sink manipulation treatment. Leaf water exchange and fruit xylem backflow were measured seven days after treatment. Leaf water status, sucrose and soluble sugar content were measured between eight to fifteen days after treatment. The same study was performed on different branches on the same trees at three dates according to fruit developmental stage: Stage I (treatment on May 14, 2024), Stage II (treatment on July 21, 2024), and Stage III (treatment on November 2, 2024) (Matias *et al*., 2023). After each source-sink manipulation treatment, any new flush was constantly removed from the treated shoots to ensure fruits were the only sink. At the end of each defoliation cycle, destructive sampling of the whole branch allowed for biomass and non-structural carbon measurements.

**Figure 1:**
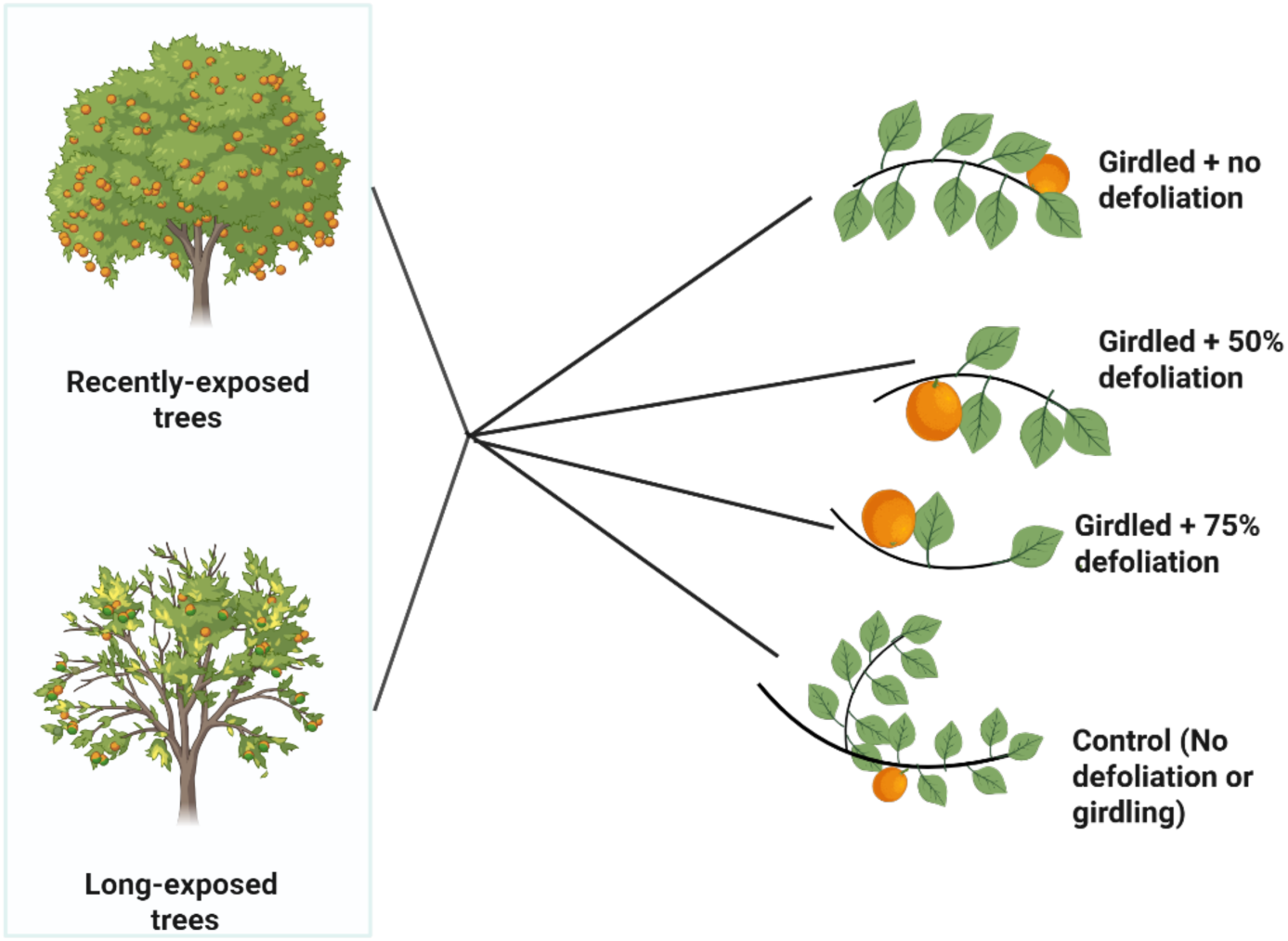
Schematic illustration of experimental conditions of source-sink manipulation (defoliation treatment) of healthy and HLB-affected field grown *Citrus* x *sinensis* ’Valencia’ trees. For each HLB disease category four branches were selected for source-sink manipulation; Control (no girdling or defoliation), Girdled (no defoliation), Girdled + 50% defoliation and Girdled +75% defoliation.

### Leaf Water Exchange

After one week of treatment, stomatal conductance (*g*_sw_) and transpiration (*E*) were measured using an porometer (LI-600; LI-COR Bioscience Inc., Lincoln, NE, USA) porometer with measurements gathered on fully developed leaves in each treatment and tree every 20 minutes from 7:00 am – 7:00 pm on a sunny day. At Stage I, sunrise and sunset were around 6:30 am and 8:10 pm respectively, at Stage II, sunrise and sunset were around 7:00 am and 8:00 pm respectively, while at Stage III, sunrise and sunset were around 7:30 am and 6:40 pm respectively. To estimate midday *g*_sw_ and *E* all measurements taken between 10:00 and 12:00 hr for midday were selected.

### Leaf Water Status

To describe the balance among leaf water content, storage, and water potential, we measured mid- day leaf water potential (Ψ_leaf_) after seven days of treatment, and leaf hydraulic properties using the bench drying pressure-volume curve method (Sack *et al*., 2010) were measured between eight to fifteen days of treatment. Well-lit leaves were collected at midday, bagged in airtight plastic bags and placed on benches for 20 minutes to allow for leaf water equilibration (Rodriguez-Dominguez *et al*., 2022). Ψ_eaf_ was measured using a Scholander pressure chamber (model 1000; PMS Instrument Co., Albany, OR, USA). The leaf weight was measured (Mettler Toledo XPE105 precision balance, Columbus, Ohio, USA), and thereafter the leaves were hydrated in water under light overnight. Next morning, the leaves were pat-dried with paper towels to remove all external water then bagged for equilibration again. After 20 minutes, the leaves were weighed and Ψ_leaf_ measured as before and then placed on bench for desiccation for 10-20 minutes, and the process was repeated again, with equilibration, weight and Ψ_leaf_ measurement and bench drying repeated seven or eight times to achieve Ψ_leaf_ less than or equal to -3 MPa, and afterwards the leaves were dried in a drying oven for 72 hours at 70°C and dry weight measured. The data collected were analyzed using a pressure-volume analysis spreadsheet tool (PVAST; Sack *et al*., 2010) from which we estimated the following variables (Also described in Table 1): turgor (pressure) potential (Ψ_p_); osmotic (solute) potential (Ψ_o_); osmotic potential at turgor loss point (π_TLP_); total and symplastic modulus of elasticity at full turgor (ɛ* and ɛ respectively); total and symplastic relative water content at turgor loss point (*RWC*\*_tlp_ and *RWC*_tlp,_ respectively); total and symplastic capacitance at full turgor (*C\**_ft_ *, C*_ft_ respectively) and at turgor loss point (*C\**_tlp_ *, C*_tlp_ respectively).

**Table 1:** List of abbreviations and variable names used in estimating gas exchange.

| Abbreviation | Variable | Units |
| --- | --- | --- |
| $\Psi_{\text{leaf}}$ | Leaf water potential | MPa |
| $\Psi_o$ | Osmotic (or solute) potential | MPa |
| $\Psi_p$ | Turgor (or pressure) potential | MPa |
| $\pi_o$ | Osmotic potential at full turgor | MPa |
| $\pi_{TLP}$ | Osmotic potential at turgor loss point | MPa |
| $\epsilon^*$ | Total modulus of elasticity at full turgor | MPa |
| $\epsilon$ | Symplastic modulus of elasticity at full turgor | MPa |
| $RWC^*_{tlp}$ | Total relative water content at turgor loss point | % |
| $RWC_{tlp}$ | Symplastic relative water content at turgor loss point | % |
| $C^*_{ft}$ | Total capacitance at full turgor | MPa<sup>-1</sup> |
| $C_{ft}$ | Symplastic capacitance at full turgor | MPa<sup>-1</sup> |
| $C^*_{tlp}$ | Total capacitance at turgor loss point | MPa<sup>-1</sup> |
| $C_{tlp}$ | Symplastic capacitance at turgor loss point | MPa<sup>-1</sup> |
| $g_{sw}$ | Stomatal conductance | mmol H<sub>2</sub>O m<sup>-2</sup> s<sup>-1</sup> |
| $E$ | Transpiration | mmol H<sub>2</sub>O m<sup>-2</sup> s<sup>-1</sup> |

### Sucrose, soluble sugar content

Leaf and fruit samples were collected after each leaf water measurement between eight and fifteen days of treatment, flash-frozen, and stored at -20°C prior to lyophilization. Non-structural carbon was determined by extracting soluble sugars, sucrose and starch from 25 mg of ground, dry leaf, stem, or fruit samples. The samples were boiled in 500 μl of ultrapure water for 10 mins at boiling point (100 °C) and agitated before centrifuging for five minutes at 13,000 rpm. After this the samples were separated into supernatants and pellets, the supernatant was used for reducing sugar and sucrose analysis.

For the sucrose analysis, 50 μl of potassium hydroxide was added to 50 μl of the supernatant and the sucrose standard solution, the samples were boiled for 10 minutes to eliminate other reducing sugars except sucrose. The samples were cooled by running under water and then 1,000 μl of 72% diluted anthrone reagent was added to each tube. The samples were agitated and incubated for 10 minutes at 100 °C and shaken again after incubation. From each sample tube, 150 μl of the incubated solution was pipetted in duplicates into a 96-well plate and absorbance was read using a spectrophotometer at 620 nm wavelength.

For the total soluble sugar analysis, 50 μl of the supernatant or the glucose standard solution were added into new tubes and 1000 μl anthrone reagent was added to each tube. The samples were agitated and incubated for 10 minutes at 100 °C and shaken after the incubation again. From each sample tube, 150 μl was pipetted in duplicates into well plate and read using spectrophotometer at 620 nm wavelength.

Linear functions based on standard curve concentrations included in each plate were generated with the standard’s absorbance to estimate reducing sugar, sucrose and starch content in the plant organs.

### Fruit diameter (Diurnal fruit shrinkage pattern)

We measured fruit diameter before defoliation treatment and at harvest to estimate fruit diameter change. The fruit diameter was measured along six equatorial transects to account for technical error especially since HLB-affected trees have lopsided fruits (Sutton *et al*., 2024). To determine the diurnal fruit shrinkage pattern and estimate xylem backflow, we measured fruit diameter at five different times of the day; predawn, mid-morning, mid-afternoon, dusk and predawn of the next day. Volume was estimated assuming a sphere according to the following:

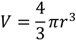

Where *V* represents the volume and r represents the mean fruit radius (diameter/2). Diurnal water loss volume was estimated as the difference between maximum fruit volume and minimum fruit volume.

### Data analysis

Data handling and statistical analysis were used R software (R Core Team, version 4.3.1, 2023). Data were analyzed using a generalized linear mixed-effects model with lmer() command from lme4 package (Bates *et al*., 2015), where tree (and fruit when analyzing fruit diurnal shrinkage pattern) was used as a random effect (nesting), and analysis of variance was performed using the anova() command. For every significant result (*P*-value < 0.05), means separation was done using the emmeans() command in the multcomp package (Hothorn *et al*., 2008). To test for relationships between variables, ggpairs() command from GGally package (Schloerke *et al*., 2025) was used to generate Pearson correlations and their corresponding *P*-values, as well as to visualize the correlations, from which linear regression models were generated for variables with significant correlations with or without disease factor using lm() command and the significant relationship was tested using anova() command at *P* value < 0.05.

## RESULTS

### 1. HLB increased leaf turgor potential

To test how HLB impacts leaf carbon-water interactions, we measured leaf sugar content, leaf hydraulic traits, diurnal and mid-day *g*_sw_ and *E* under varying source:sink ratios. We did not find any significant effect of source-sink manipulation on leaf sucrose or soluble sugar (Fig. 2, Supplemental Table 2). However, exposure to HLB, fruit development stage and their interaction affected leaf sucrose and total soluble sugar (for all effects, *P* < 0.0001, Fig. 2A; Supplemental Table S2). Leaf sucrose content of recently-exposed trees remained lower than long-exposed trees except at Stage II where both HLB disease categories were similar. Leaf total soluble sugar was higher in long-exposed trees at Stage I, than all other disease and fruit development stage combinations, which did not differ from each other (Fig. 2B).

**Figure 2:**
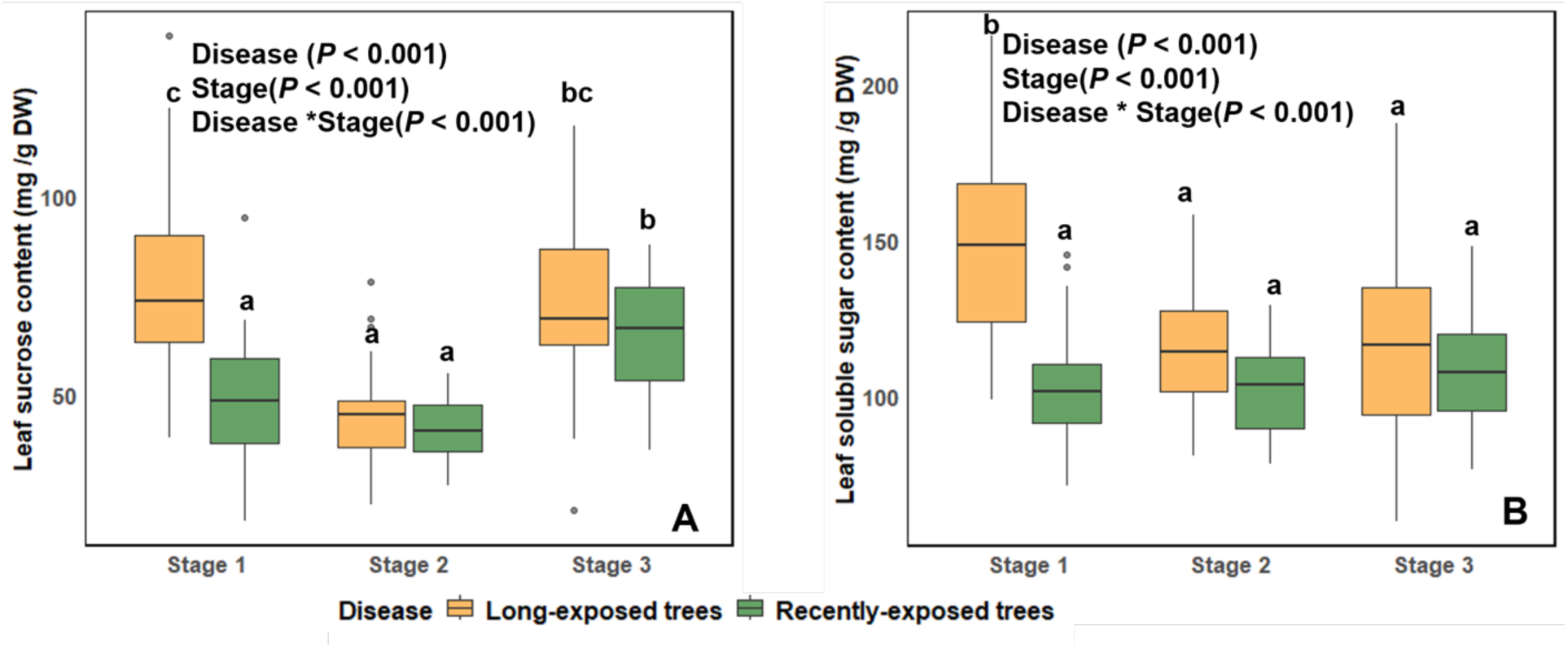
Leaf sucrose content (A) and total soluble sugar (B) of field-grown *Citrus* x *sinensis* ’Valencia’ trees in a study of huanglongbing disease and source-sink manipulation. Data collected during Stage I, II and III of fruit development. Groups labeled with different letters are significantly different (Tukey’s Honestly Significant Difference; *P* < 0.05).

We observed distinct leaf hydraulic traits depending on exposure to HLB, fruit development stage, and source-sink manipulation (Fig. 3 and 4; Supplemental Table 3). ɛ*, ɛ, *C*_ft_, *RWC*_tlp_, *C*_tlp_, π_TLP_, π_o_, ψ_o_ and ψ_p_ were affected by the interaction of HLB exposure with fruit development stage (Fig. 3). *RWC\**_tlp_ was the hydraulic traits that was only affected by HLB disease exposure duration. We observed that ψ_p_, ɛ*,ɛ, *RWC*_tlp_, and *C*_tlp_ had similar patterns, in which long-exposed trees had lower leaf hydraulic traits at Stages I and III, but increased at Stage II, while recently-exposed trees responded in the opposite dynamic. However, *C*_f_, π_o_, and ψ_o_ were higher in long- exposed trees at Stages I and III, but low at Stages II compared to recently-exposed trees. Furthermore, although ψ_o_, ψ_p_ , π_TLP_, and π_o_, showed significant interactions between fruit development stage and exposure to HLB (*P* = 0.01, Fig. 3A; *P* = 0.04, Fig. 3B; *P* = 0.006, Fig. 3C and P =0.004, Fig. 3D, respectively), pairwise comparisons did not reveal separation among category means, indicating that the interaction effect is due to differing dynamics rather than large contrasts between any particular categories. We also observed that long-exposed trees had lower *RWC\**_tlp_ than recently-exposed trees.

**Figure 3:**
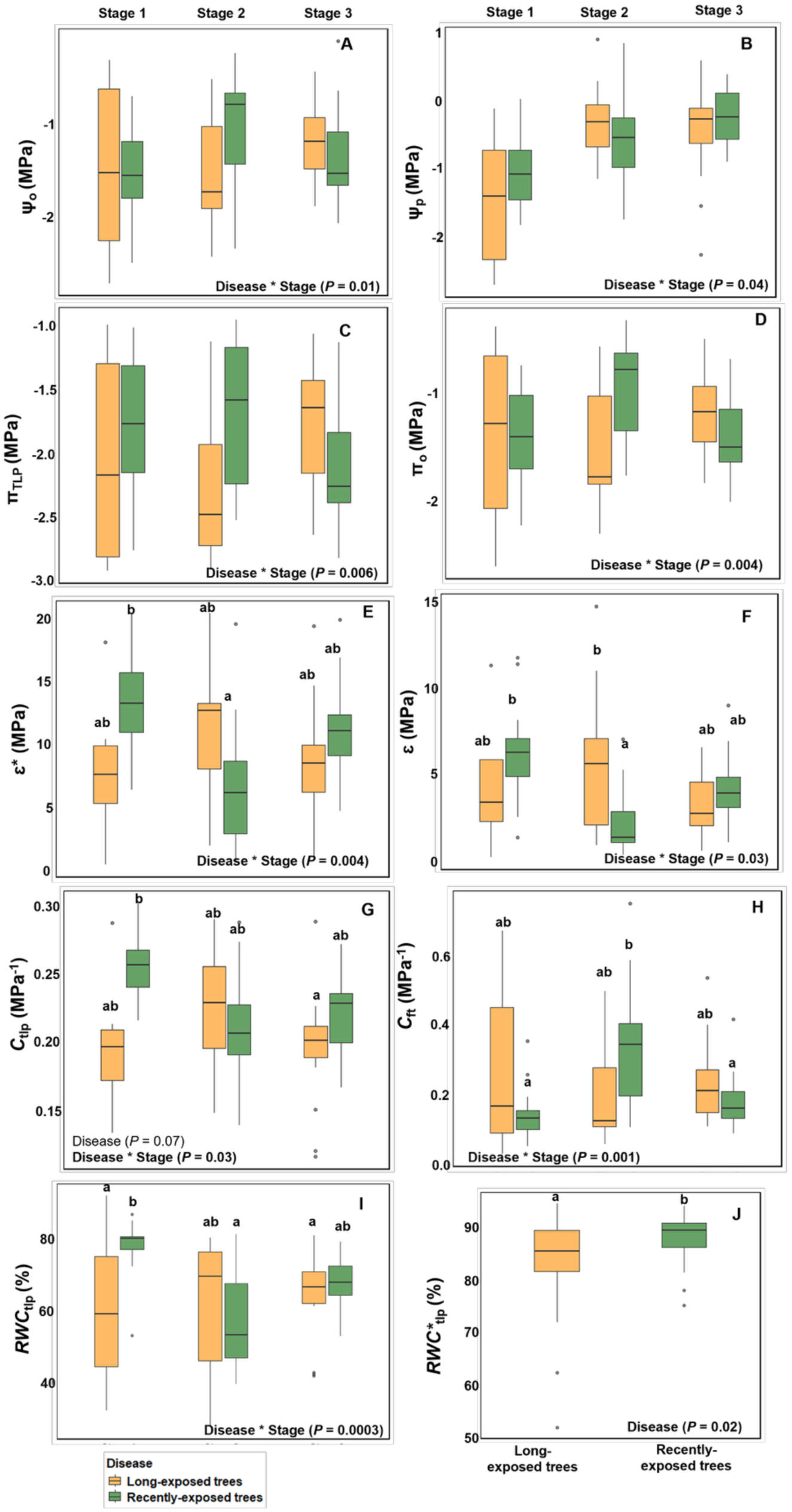
Effect of huanglongbing disease exposure duration on leaf hydraulic properties of field- grown *Citrus* x *sinensis* ’Valencia’ trees in a study of disease and source-sink manipulation at Stages I, II and III of fruit development. Hydraulic variabls include osmotic potential (A; Ψ_o_ ); pressure (turgor) potential (B; Ψ_p_ ); osmotic potential at turgor loss point (C; π_TLP_); osmotic potential at full turgor (D; π_o_); symplastic relative water content at turgor loss point (E; *RWC*_tlp_); total relative water content at turgor loss point (F; *RWC*_tlp_); total modulus of elasticity at full turgor (E; ɛ*); symplastic modulus of elasticity at full turgor (F; ɛ); symplastic capacitance at turgor loss point (G; *C*_tlp_) and symplastic capacitance at full turgor (H; *C*_ft_). Groups labeled with different letters are significantly different (Tukey’s Honestly Significant Difference; *P* < 0.05).

**Figure 4.**
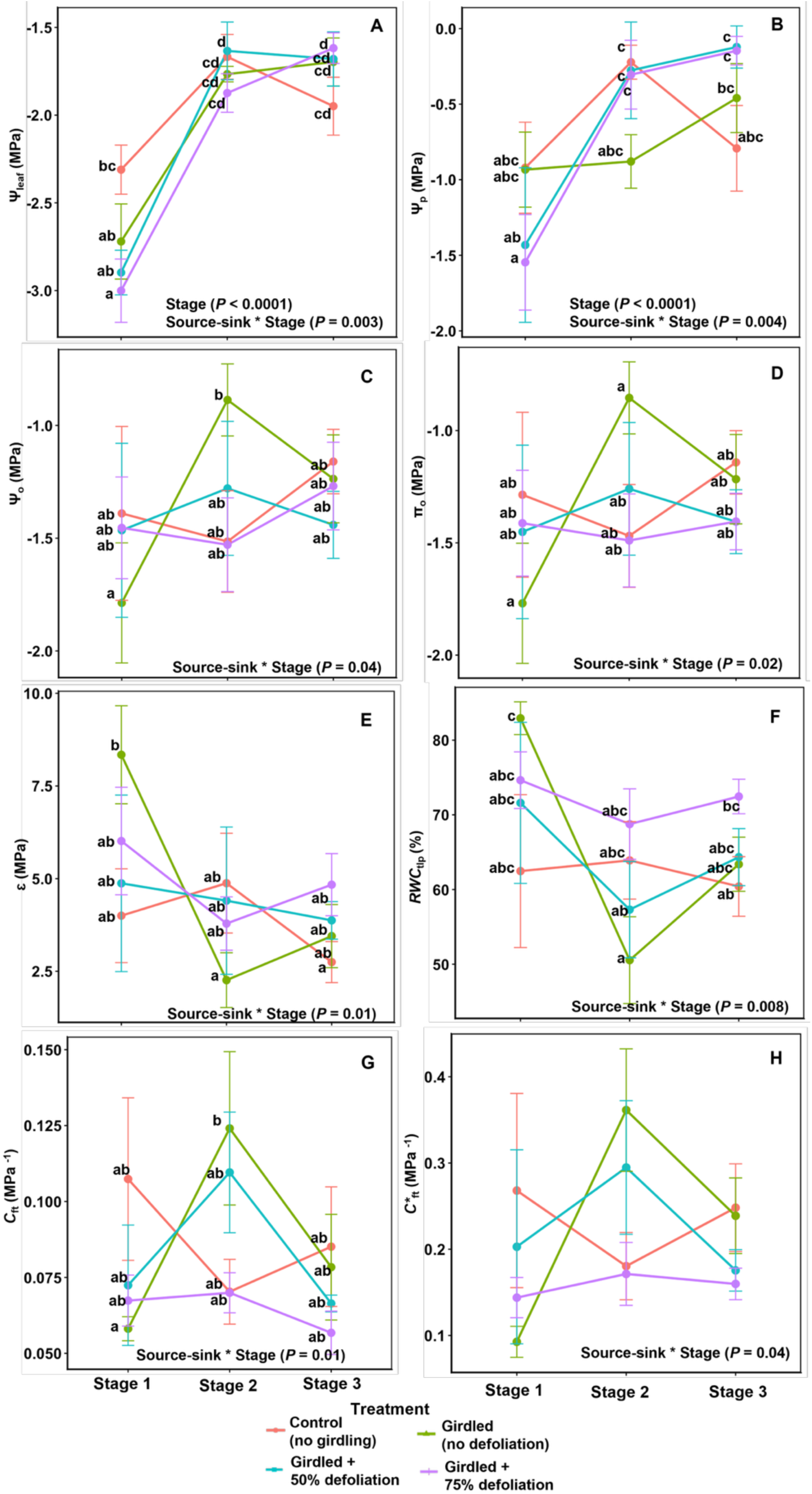
Effect of source-sink manipulation on leaf hydraulic properties of field-grown *Citrus* x *sinensis* ’Valencia’ trees in a study of huanglongbing disease and branch-level source-sink manipulation measured during Stage I, II and III of fruit development. Variables include leaf water potential (A; Ψ_leaf_); pressure (turgor) potential (B; Ψ_p_); osmotic potential (C; Ψ_o_); osmotic potential at full turgor (D; π_o_); symplastic modulus of elasticity at full turgor (E; ɛ); symplastic relative water content at turgor loss point (F; *RWC*_tlp_); symplastic capacitance at full turgor (G; *C*_ft_) and capacitance at full turgor (H; *C*_ft_). C denotes control (no girdling or defoliation), G + 0 denotes girdled (no defoliation); G + 50 denotes girdled + 50% defoliation; G + 75 denotes girdled + 75% defoliation treatments. Groups labeled with different letters are significantly different (Tukey’s Honestly Significant Difference; *P* < 0.05).

The interaction of source-sink manipulation and fruit development stages affected ψ_leaf_, ψ_p_, ψ_o_, π_o_, ɛ, *RWC*_tlp_, *C*_f_ and *C\**_ft_ (Fig. 4). ψ_leaf_ had the most pronounced effect at Stage I of fruit development, where both girdling and defoliation reduced ψ_leaf_. However, 50% defoliation at Stage II and 75% defoliation at Stage III increased ψ_leaf_. Defoliation induced contrasting effects on ψ_p_ across the fruit developmental stages; defoliation reduced ψ_p_ at Stage I, but increased it at Stages II and III. ψ_o_, π_o_, ɛ, *RWC*_tlp_, and *C*_ft_ were only affected by girdling; girdling decreased ψ_o_, π_o_, and *C*_f_ at Stage I and increased at Stage II. On the other hand, girdling increased ɛ and *RWC*_tlp_ at Stage I but decreased them at Stage II.

Source-sink manipulation did not affect *g*_sw_. However, there was possible effect on *E* (Fig. 5; Supplemental Table 4). HLB exposure, fruit development stage and their interaction affected *g*_sw_ (Fig. 5A). At Stages I and III, leaves of long-exposed trees had lower *g*_sw_. However at Stage II, *g*_sw_ was slightly higher in long-exposed trees. The effect of source-sink manipulation on diurnal *g*_sw_ differed in both HLB disease categories (Supplemental Fig. S1A). In leaves of long-exposed trees, *g*_sw_ in defoliated branches peaked earlier than non-defoliated branches, and leaves of 75% defoliated had higher *g*_sw_ which remained consistently higher over the course of the day. Similar to long-exposed trees, *g*_sw_ of defoliated, recently-exposed trees peaked earlier, however the non- defoliated branches maintained higher *g*_sw_ throughout the day. Similarly, *E* was affected by HLB disease exposure and fruit development stage and their interaction, where long-exposed trees had lower *E* than recently-exposed trees at Stages I and III of fruit development (Fig. 5B). We observed a possible increase in *E* in defoliated branches. Leaf *E* to source-sink manipulation also differ in the two HLB disease categories studied. In leaves of long-exposed trees, 75% defoliated and girdled non-defoliated shoots transpired more than other categories, while in recently- exposed trees, the non-defoliated branches transpired more diurnally (Supplemental Fig. S1B).

**Figure 5.**
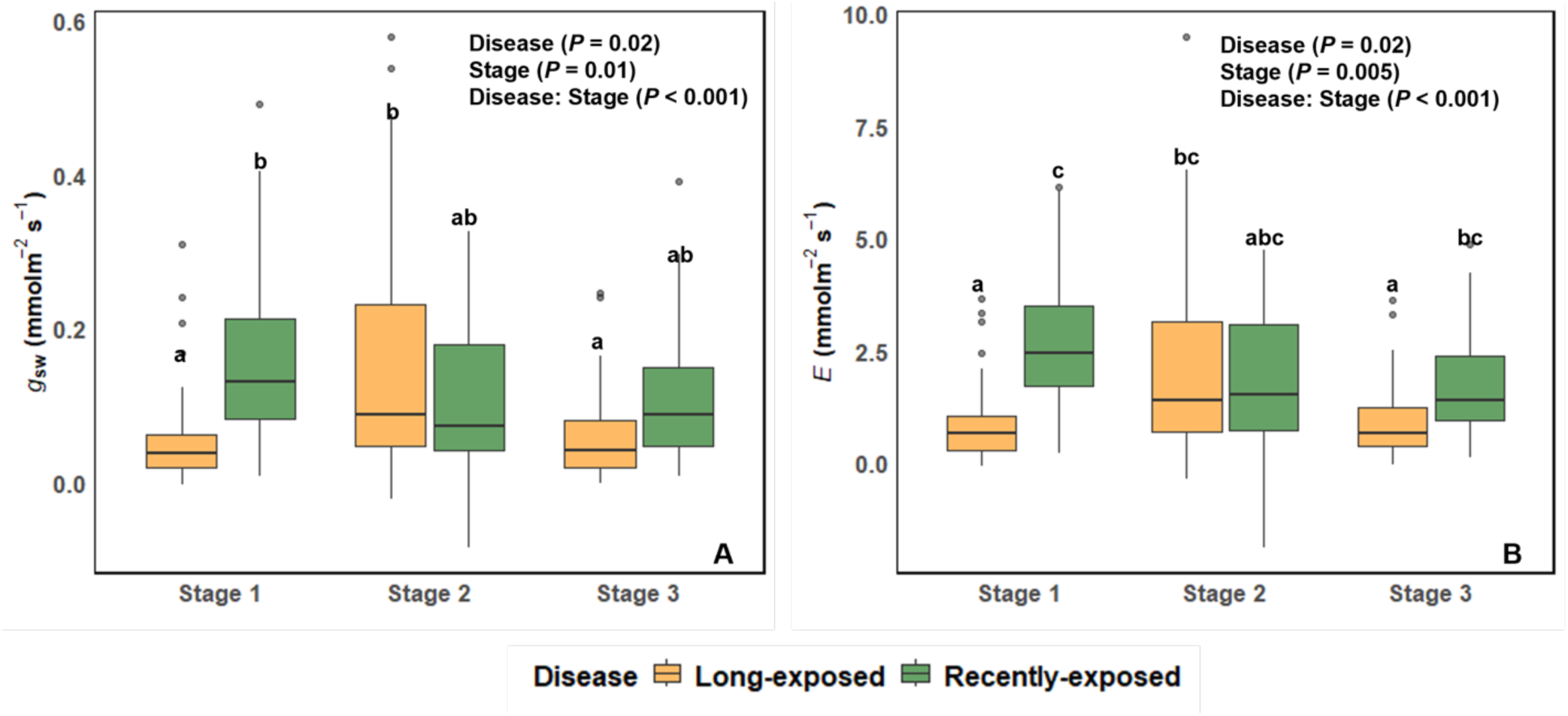
Midday leaf stomatal conductance (*g*_sw_; A) and transpiration (*E*; B) of field-grown *Citrus* x *sinensis* ’Valencia’ trees in a study of huanglongbing disease exposure and various source-sink manipulations measured during Stages I, II and III of fruit development. Groups labeled with different letters are significantly different (Tukey’s Honestly Significant Difference; *P* < 0.05).

### HLB increased xylem backflow which was further exacerbated by defoliation

To assess the impact of varying source:sink ratio on fruit water dynamics, we measured fruit sucrose, total soluble sugars and changes in diurnal fruit diameter. We found no significant effect of source-sink manipulation on fruit sugar content (Fig. 6, Supplemental Table 5). The interaction of fruit development stages and HLB exposure impacted fruit sucrose (Fig. 6A). Fruit development stage was by far the larger effect: fruit sucrose increased as fruit developed regardless of the disease categories. Fruit soluble sugar in both HLB disease categories was similar in Stages I and III, while at Stage II fruit of long-exposed trees had lower soluble sugar (Fig. 6B).

**Figure 6.**
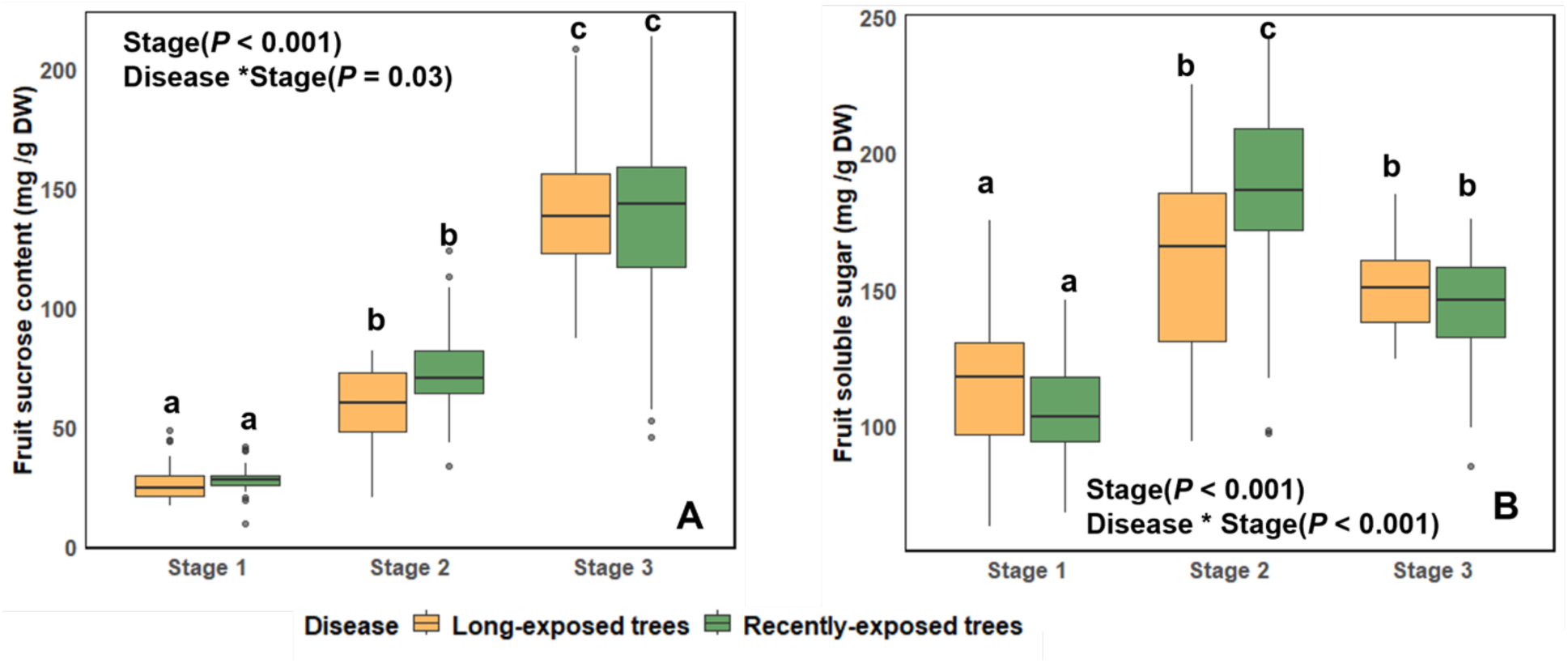
Fruit sucrose content (A) and soluble sugar (B) of field-grown *Citrus* x *sinensis* ’Valencia’ trees in a study of huanglongbing disease exposure and various source-sink manipulations at Stages I, II and III of fruit development. Groups labeled with different letters are significantly different (Tukey’s Honestly Significant Difference; *P* < 0.05).

HLB disease exposure changed the pattern of fruit shrinkage response to source-sink manipulation across the three stages studied (Fig. 7; Supplemental Table 6). At Stages I and II (Fig. 7A, B), regardless of disease exposure duration, maximum diurnal shrinkage (MDS) occurred mid-afternoon, in all source-sink manipulation treatments, except in 75% defoliated shoots where maximum shrinkage occurred by mid-morning, while the other source-sink manipulation treatments differed only in magnitude but had the same diurnal pattern. The fruit shrinkage pattern of 75% defoliated shoots was distinct in that these fruits shrank less or earlier than other treatments in all combinations of fruit development stage and disease exposure. At Stage I, fruits of long-exposed trees had greater shrinkage than fruits of recently-exposed trees while at Stage II, the long-exposed trees shrank slightly more. At Stage III (Fig. 7C), we observed a consistent pattern across treatments. Fruits expanded between pre-dawn and mid-morning, and then shrank from mid-morning to mid-afternoon or dawn depending on HLB exposure duration and or source-sink manipulation. Similar to Stages I and II, MDS occurred mainly at dusk in long-exposed trees at Stage III, except for 75% defoliated branches which occurred much earlier by mid-afternoon. Contrastingly, fruits of recently-exposed shoots reached MDS at mid-afternoon except for 50% defoliated branches where MDS occurred at dusk. Similar to what we observed at Stages I and II, fruits of 75% defoliated branches shrank less than other treatments.

**Figure 7.**
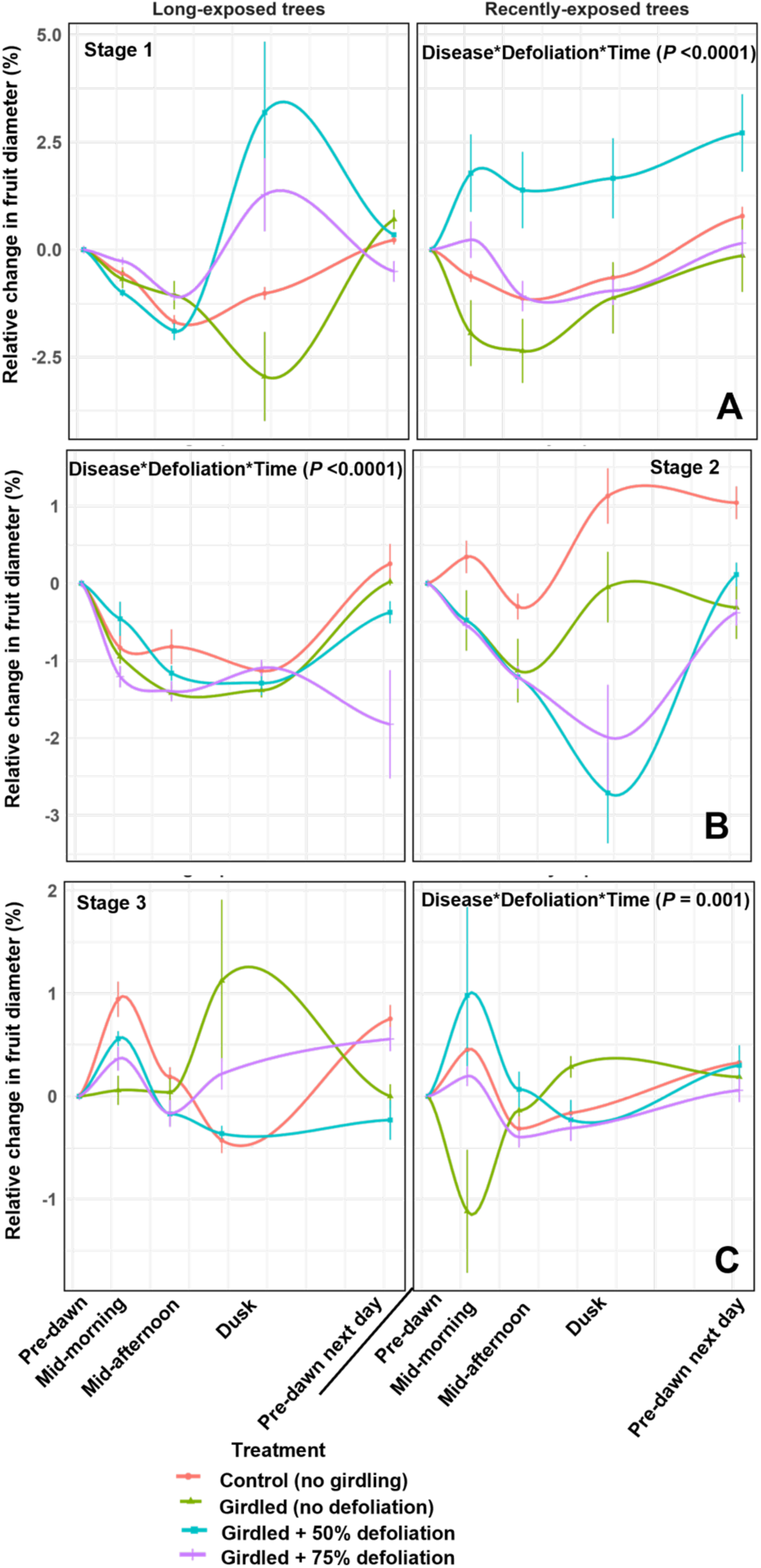
Fruit diurnal shrinkage in field-grown *Citrus* x *sinensis* ’Valencia’ trees in a study of huanglongbing (HLB) disease and various source-sink manipulations. Data collected during Stage I (A), II (B) and III (C) of fruit development. Error bars represent standard error (n = 10)

Only HLB affected both total volume of fruit water loss and relative fruit water loss (Fig. 8; Supplemental Table S7), although total fruit water loss was only affected at Stage I of fruit development, and relative fruit water loss was affected at Stages II and III, in which long- exposed trees consistently had greater relative fruit water loss than recently-exposed trees. Relative fruit water loss was not different in Stage I, but total fruit water loss was greater in long- exposed trees at Stage I.

**Figure 8.**
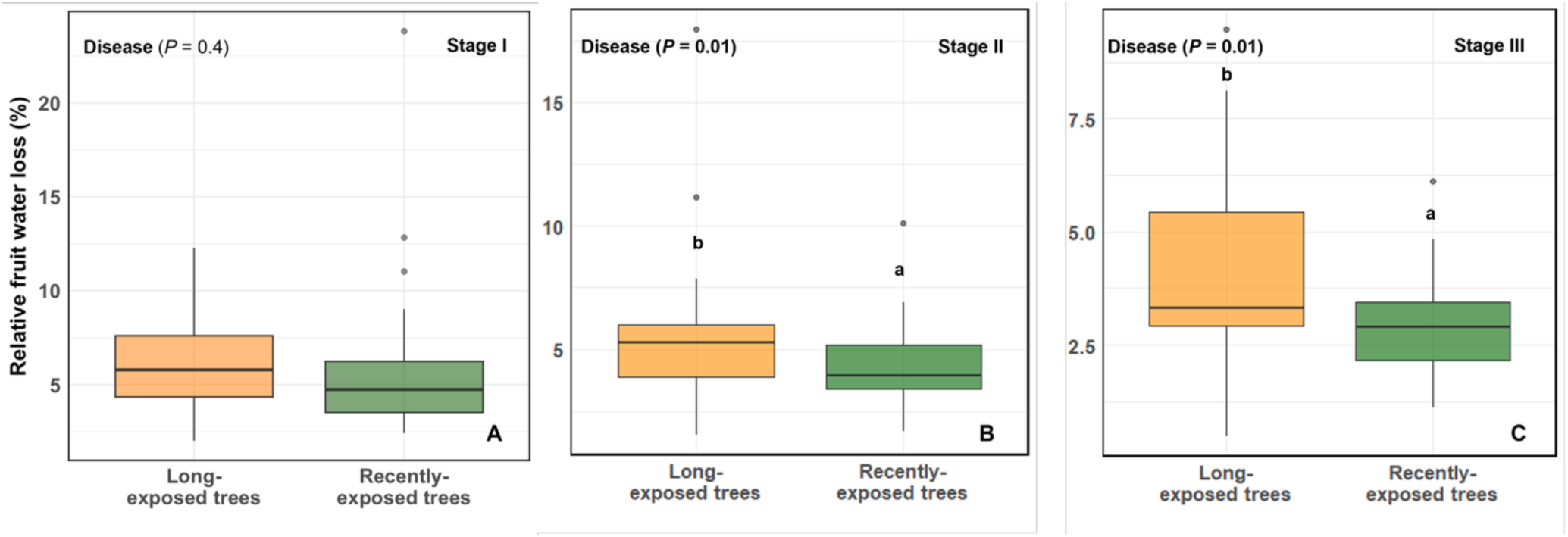
Effect of huanglongbing (HLB) exposure duration on fruit daily water loss (% of total maximal water content) in fruit development Stages I, II, and III (A, B, and C, respectively) of field grown *Citrus* x *sinensis* ’Valencia’ trees in a study of HLB and various source-sink manipulations. Groups labeled with different letters are significantly different (Tukey’s Honestly Significant Difference; *P* < 0.05).

To assess the relationship between source-sink dynamics and water relations, we explored correlations among leaf hydraulic traits, leaf fruit water content, fruit sucrose and soluble sugar content (Supplemental Fig. S2, Supplemental Table S9). We observed that total fruit water loss correlated with ψ_p_, π_o_, ψ_leaf_, fruit soluble sugar and fruit sucrose content. Interestingly, we found leaf sucrose content only correlated with *C*_tlp_. Leaf *E* positively correlated with ɛ, ɛ*, *RWC*_tlp_, and negatively correlated with *C\**_ft_, *C*_tlp_, π_o_, and ψ_o_. Similarly, *g*_sw_ correlated positively with *E*, ψ_p_, ɛ, ɛ*, *RWC*_tlp_, *C\**_tlp_, and *C*_tlp_, negatively with *C\**_ft_, π_o_, and ψ_o_. Meanwhile, fruit sucrose content correlated positively with ψ_leaf_ negatively in Stage I and positively in Stages II and III. Fruit sucrose also correlated negatively with *C*_tlp_, and *C\**_tlp_ across all the stages. We only observed a relationship between fruit sucrose and ɛ at Stage III. Fruit soluble sugar also correlated with ɛ and ψ_leaf_ (Supplemental Fig. S2). Linear regression analysis further confirmed the positive relationships of *g*_sw_ with ψ_p_, ɛ, and *RWC*_tlp_, and negative relationships of *g*_sw_ with *C*_ft_, π_o_, and ψ_o_ (Fig. 9). In addition, both ψ_leaf_ and ψ_o_ increased with increasing fruit water content (Fig. 10; Supplemental Table S9).

**Figure 9.**
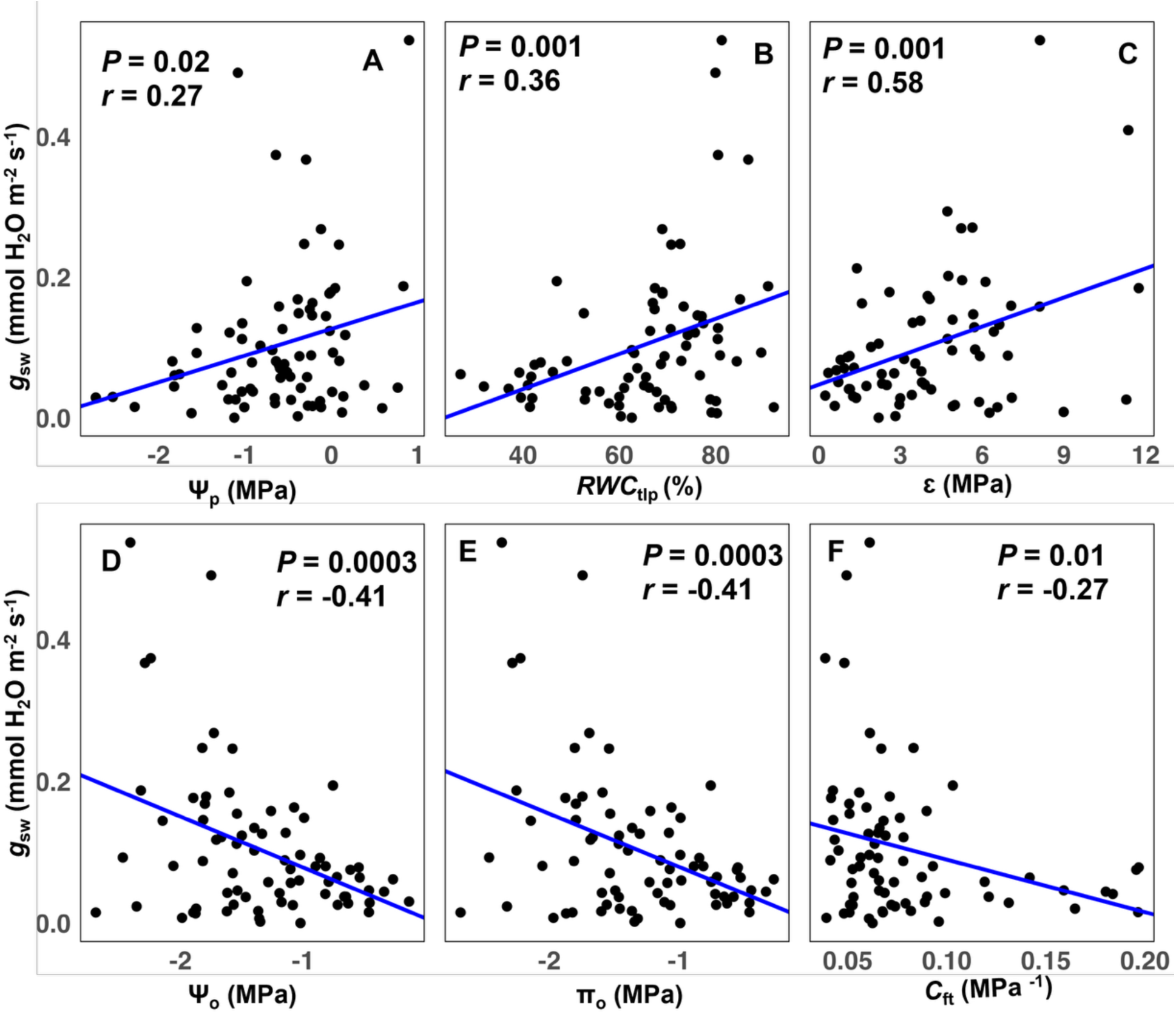
Midday leaf stomatal conductance (*g*_sw_) relationship with some leaf hydraulic traits of field-grown *Citrus* x *sinensis* ’Valencia’ trees in a study of huanglongbing disease exposure and various source-sink manipulations measured during Stages I, II and III of fruit development. A, Ψ_p_ - Pressure (turgor) potential; B, *RWC*_tlp_ -symplastic relative water content at turgor loss point; C, ɛ - symplastic modulus of elasticity at full turgor; D, Ψ_o_ - osmotic potential; E, π_o_ - osmotic potential at full turgor; F, *C*_ft_ - symplastic capacitance at full turgor.

**Figure 10.**
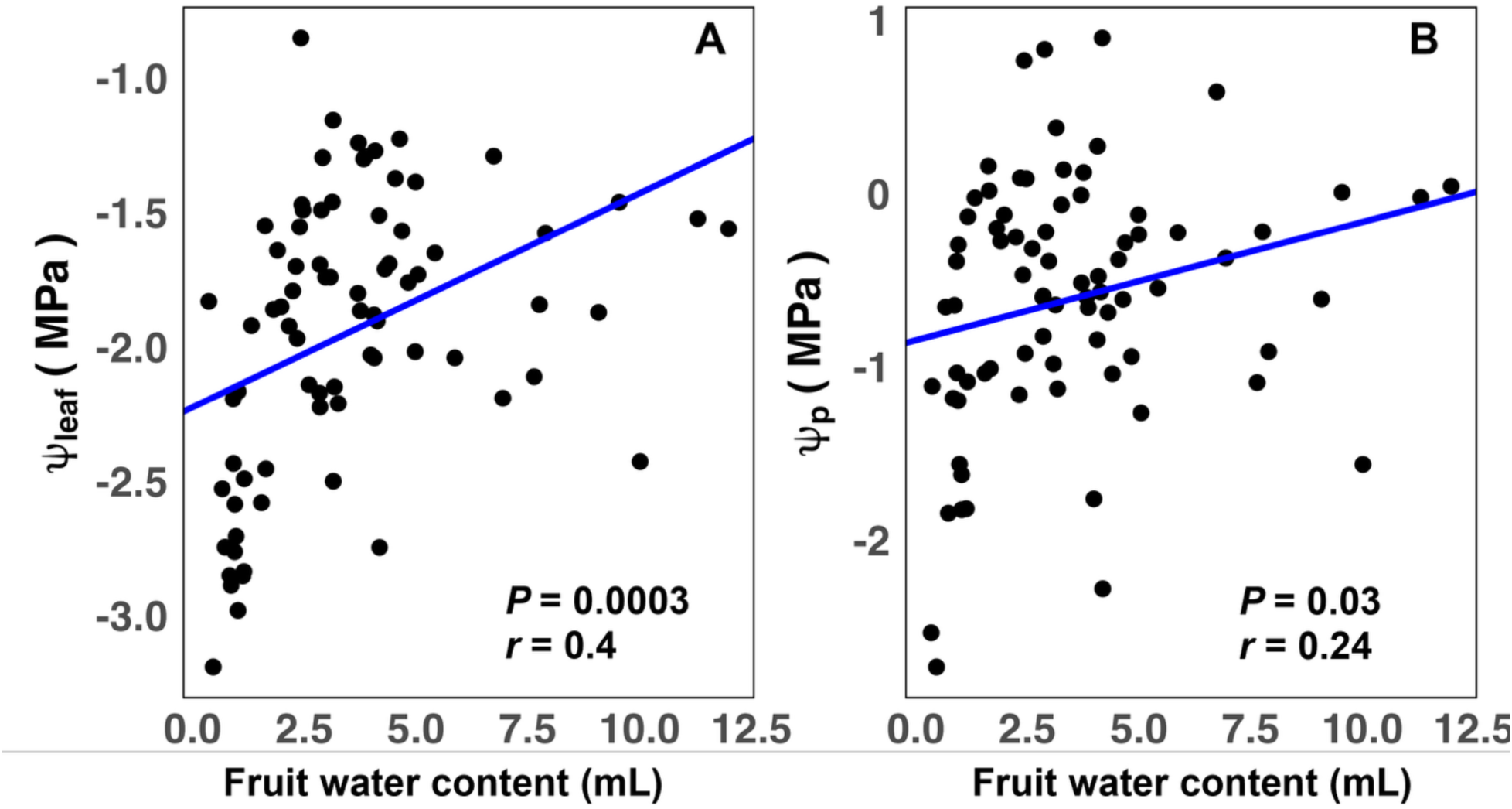
leaf water potential (Ψ_leaf_) and turgor potential (Ψ_p_) relationship with fruit water content of field-grown *Citrus* x *sinensis* ’Valencia’ trees in a study of huanglongbing disease exposure and various source-sink manipulations measured during Stages I, II and III of fruit development.

## DISCUSSION

In this study, we investigated how source-sink and water dynamics interact as HLB develops over time. HLB causes water deficit in HLB-affected plants, due to reduced root growth caused by phloem blockage, hypothetically disrupting the relationship between carbohydrate and water transport (Johnson *et al*., 2014; Sutton *et al*., 2025). This is the first study to address how citrus trees acclimate to disease-induced water deficits and the first to address the impacts of source:sink on hydraulic acclimation. Prolonged HLB exposure appears to induce acclimation to water deficits, by altering leaf hydraulic traits (like leaf capacitance, modulus of elasticity) and increasing fruit xylem backflow. However, this acclimation response is limited when sink activity is high, likely due to tradeoffs between carbon transport for growth and carbohydrate accumulation for osmotic adjustment (Fig. 11).

**Figure 11:**
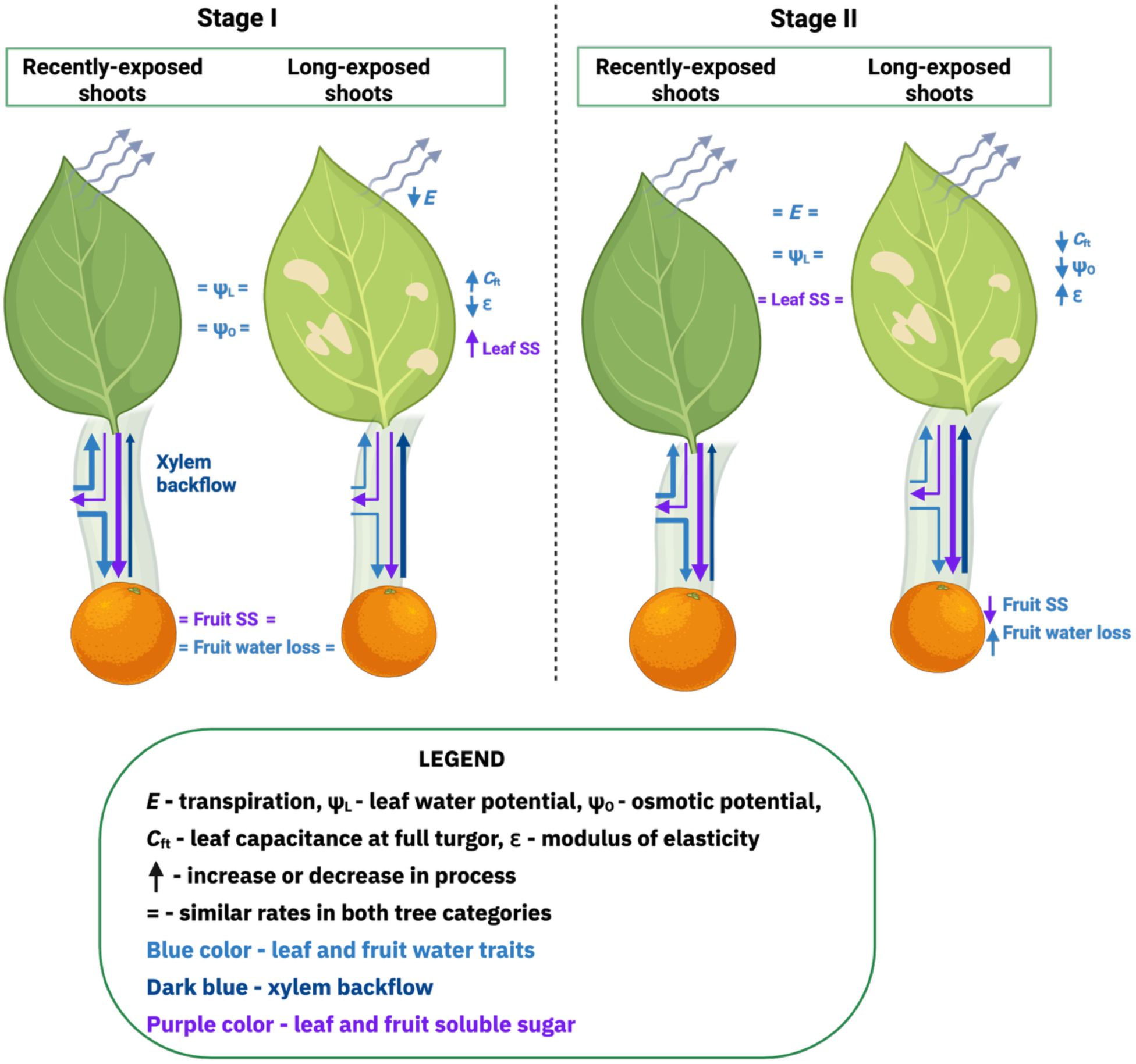
Graphical illustration of plant water-carbon dynamics in long-exposed and recently-exposed HLB trees during Stages I and II of fruit development. During early Stage I, long- exposed HLB-affected shoots exhibited reduced transpiration, modulus of elasticity, increased capacitance and increased fruit xylem backflow, suggesting acclimation to water deficit under prolonged HLB exposure. In stage II, however, these acclimations were less pronounced reflecting increased carbon transport to rapidly growing fruit during this stage.

We hypothesized that HLB alters leaf turgor pressure (ψ_p_) in leaves of affected trees, and to our knowledge this is the first study to directly examine this relationship. Several studies have reported that HLB reduces stem sap flow and decreases leaf and xylem water potential (Hamido *et al*., 2019; Kwakye and Kadyampakeni, 2023), which would diminish ψ_p_. However, previous work showed that HLB also increases nonstructural carbohydrate (sucrose, starch) in leaves of HLB-affected trees (Cai *et al*., 2022), which could increase ψ_p_ by increasing ψ_o_. The present work sought to quantify the net effects of these countervailing disease impacts to ascertain whether HLB impairs leaf capacity to draw water from the xylem and maintain turgor pressure. The results suggest that, while HLB affects all components of turgor, the responses interact dynamically with fruit development.

Leaf ψ_o_ is one of the determinants of turgor pressure, because increased solute concentration reduces ψ_o_, promoting water influx and increasing turgor pressure to support carbon transport (Zhang *et al*., 2024). In the present study, leaves of long-exposed HLB-affected trees accumulated more sucrose and soluble sugars, particularly at Stage I, which we expected would translate to decreased (more negative) ψ_o_. However, at Stage I, ψ_o_ was slightly less negative in leaves of long-exposed trees relative to recently-exposed trees. At Stage II, when soluble sugars and sucrose were lower and similar across the two HLB categories, ψ_o_ tended to become more negative in long-exposed trees. Other sugar accumulation, dehydration and osmolyte accumulation could also influence solute potential (Gersony *et al.,* 2020; Babst *et al*., 2022). Together, these results suggest that sucrose accumulation in long-exposed trees may contribute to osmotic adjustment and acclimation to disease-induced water deficit, although this response is fruit stage-dependent.

As hypothesized, HLB exposure duration affected ψ_p_, particularly at Stage I and II of fruit development, however, the lack of statistical separation among means warrants cautious interpretation. This lack of clear differences reflects the inherently low operating turgor of citrus (0.5 MPa; Walker *et al*., 1983) compared with other species, such as apple (0.95–1.70 MPa; Davies and Lakso, 1979) and red oak (0.8–1.6 MPa; Gersony *et al*., 2020). This low turgor may limit detectable shifts in ψ_p_. Additionally, because all trees in this study were HLB-exposed, differences between disease categories may have been constrained. Moreover, because citrus trees exhibit isohydric behavior (Romero-Trigueros *et al*., 2021), characterized by reduced stomatal conductance under water deficit to conserve water, as observed in leaves of HLB-affected trees, this regulation likely helps maintain ψ_leaf_, which might limit effects on turgor. From the present study we also observed that ψ_p_ positively correlated with ψ_leaf_ and *g*_sw_, reinforcing the role of stomata in maintaining ψ_p_. Further study including healthy controls could clarify the HLB disease effects on ψ_p_.

In addition to the more immediate impacts on ψ_leaf_, ψ_p_, and ψ_o_, HLB disease exposure also impacted longer term leaf hydraulic traits including elasticity (ɛ), capacitance (*C*), osmotic potential at turgor loss point (π_TLP_), and relative water content at turgor loss point (*RWC*_tlp_). In the present study, long-term exposure to HLB was associated with more negative π_TLP_, reduced *RWC*_tlp_, less ɛ* and ɛ, and increased *C*_ft_, all of which are traits linked to drought acclimation. π_tlp_ is often recognized as a key predictor of drought tolerance because a more negative value extends the range of ψ_leaf_ over which leaves remain turgid (Bartlett *et al*., 2012), further confirmed by lower *RWC*_tlp_. *RWC*_tlp_ has also been linked to maintenance of stomatal conductance, hydraulic conductance, photosynthetic activity and even growth under low soil moisture (Hochberg *et al*., 2017). Thus, the lower the *RWC*_tlp_, the more acclimated the leaves are to water deficit (Sack *et al*., 2003; Hochberg *et al*., 2017). Reduced cell wall elasticity (lower ɛ) can facilitate maintenance of ψ_p_, while higher capacitance enhances the leaf capacity to release stored water and buffer declines in ψ_leaf_ (Touchette *et al*., 2014; Xiong and Nadal, 2020; Huo *et al*., 2022). These coordinated adjustments suggest that long-term HLB exposure induces acclimation to water deficit, consistent with reports of improved stomatal conductance and transpiration in long-exposed trees relative to more recently-exposed trees (Keeley *et al*., 2022). These structural adjustments may also explain the limited effect of HLB on ψ_p_.

However, there may also be tradeoffs between source-sink transport and leaf water-deficit acclimation. The acclimation effects were reversed temporarily during Stage II, a period of high sink activity. At this stage, foliar sucrose declined, indicating enhanced carbon export under strong sink demand, despite continued low *g*_sw_, suggesting that the water deficit conditions continued. Although ψ_o_ was more negative at Stage II, ɛ and *RWC*_tlp_ increased and *C*_ft_ reduced, all of which suggest reduced acclimation effect by increasing cell rigidity, reduced hydraulic buffering capacity from stored water within the plant and earlier turgor loss at relatively higher water contents. These shifts suggest a tradeoff between maintaining water-deficit acclimation and supporting carbon exports when sink demand is high. Further evidence for this tradeoff emerged from source-sink manipulations. Leaf hydraulic trait responses to shoot girdling varied by fruit developmental stages. At Stage II, girdled shoots showed acclimation response by decreasing *RWC\**_tlp_ and ɛ, as well as increasing *C*_ft_. In contrast, defoliated shoots lacked these acclimation responses, which further reinforces that that high sink demand can attenuate water-deficit acclimation mechanisms.

Long-term exposure to HLB reduced *g*_sw_ and *E*. However, increasing sink demand attenuated these negative effects. Across stages, long-exposed trees exhibited lower *g*_sw_ and *E*, with only a slight increase in *g*_sw_ at Stage II, indicating sustained water deficit responses. This contrasts with reports by Keeley *et al*. (2022), who observed increased *g*_sw_ with prolonged HLB exposure, though that study was performed under controlled conditions on non-fruiting trees, conditions whose relevance is highlighted by the present work. Diurnal patterns of *g*_sw_ and *E* at Stage II also revealed that low source:sink ratio induced early and greater stomatal activity which oscillated diurnally in long-exposed trees, but not in recently-exposed trees. Moreover, leaf *E* and *g*_sw_ were negatively correlated with ψ_o_, suggesting that greater osmotic adjustment enhanced *g*_sw_ and maintained *E. g*_sw_ positively correlated with *RWC*_tlp_ and ɛ, suggesting that increased *g*_sw_ was associated with increased cell rigidity and earlier turgor loss. Conversely, *g*_sw_ was negatively correlated with *C*_ft_, which indicates that lower *g*_sw_ coincided with greater hydraulic buffering capacity. Thus, long-exposed tree acclimation to water deficit is modulated by sink activity and may be temporarily overridden during periods of high carbon export demand, revealing a dynamic tradeoff between hydraulic stability and carbon allocation in HLB-affected trees. In this dynamic, *g*_sw_ responds to the other hydraulic variables, maintaining ψ_leaf_ and mitigating potential deficits.

Xylem backflow from fruits occurs throughout the growing season and has been reported in grapes, kiwifruit, plum, apple, sweet cherry and citrus (Elfving and Kaufmann, 1972; Lang, 1990; Greenspan *et al*., 1994; Morandi *et al*., 2010; Brüggenwirth *et al*., 2016; Winkler and Knoche, 2021). Several factors influence this process, including fruit developmental stage, time of day, transpiration rate, and atmospheric vapor pressure deficit (Winkler and Knoche, 2021). In grape, fruit xylem backflow plays an important role in discharging excess water to enable the onset of ripening (Zhang and Keller, 2017), while in citrus, xylem backflow has been associated with water recycling when transpiration demand exceeds water supply and is closely linked to source-sink carbon transport (Goldschmidt and Koch, 1996). When transpiration demand exceeds water supply to the leaf, a steep water potential gradient may drive xylem backflow from fruit to leaves, recycling water to maintain leaf turgor, but reducing fruit turgor, and thereby supporting phloem pressure gradients necessary for continued carbon transport.

HLB appears to increase fruit xylem backflow as a compensatory mechanism to maintain leaf hydraulic function under disease-induced leaf water deficit. Previous studies demonstrated that HLB predisposes trees to water deficit, due to carbon starvation in roots and declining root activity which reduces water uptake and availability (Johnson *et al*., 2014; Hamido *et al*., 2017; da Silva *et al*., 2021). In the present study, we observed that long-term HLB exposure increased fruit maximum daily shrinkage (MDS), indicating a greater reliance on fruit xylem backflow as a compensatory response to leaf water deficit.

Xylem backflow impacts on fruit water reduced as fruit developed. Although Stage I is characterized by low sink demand with low sucrose levels, we observed that this stage exhibited the greatest proportional fruit shrinkage (Fig. 6), which progressively declined as fruit developed. As fruit sucrose content increased with fruit development, this pattern suggests enhanced osmoregulation and turgor maintenance in fruits thereby increasing water retention and reducing fruit shrinkage. This pattern is consistent with previous reports describing the difficulty of maintaining high turgor in rapidly expanding citrus fruit sinks (Huang *et al.,* 1992; Goldschmidt and Koch, 1996). We also observed a positive correlation between total fruit water loss and leaf ψ_p_. The relatively high xylem backflow during Stage I may also explain the less negative leaf ψ_o_ observed, particularly in long-exposed trees, despite their higher leaf sucrose content. At Stage III, fruit shrinkage exhibited a distinct pattern compared with earlier stages, with fruit expansion occurring between pre-dawn and mid-morning. This may be attributable to environmental conditions during that period, particularly lower temperatures relative to Stages I and II, which likely reduced vapor pressure deficit and consequently transpiration. Thus, xylem backflow is closely linked with leaf turgor, and this relationship is influenced by fruit developmental stages and environmental conditions.

Similar to leaves, fruits exhibited a water-carbon tradeoff under water deficit conditions. Fruits with lower proportional shrinkage showed greater increase in diameter compared with those exhibiting higher MDS. Fruit water loss was positively correlated with leaf turgor and fruit sugars, suggesting that increased leaf turgor may reduce fruit turgor to maintain the phloem pressure gradient and maintain fruit growth (Goldschmidt & Koch 1996). Despite continued carbon import, greater fruit water loss may have limited cell expansion, indicating that fruit growth under water deficit was constrained by both carbon availability and water retention. Girdling increased diurnal fruit growth in recently-exposed trees at Stage I, likely due to carbohydrate accumulation in leaves and subsequent mobilization to active sinks. However, this response was not observed in long-exposed trees, suggesting greater impairment of phloem transport in the long-exposed group. Fruits on 75% defoliated shoots showed a distinct pattern, particularly in long-exposed trees, characterized by earlier MDS or reduced shrinkage, which did not translate to increased diurnal fruit growth. Although defoliation reduced transpiration rate per shoot by reducing transpirational surface area, carbon supply was also limited and thereby reduced fruit growth. This pattern demonstrates the importance of both carbohydrate and water supply to both leaves and fruits. Reducing water and carbon supply induces a tradeoff between allocation of both water and carbohydrates among both organs.

Carbon delivery to fruits and sucrose accumulation in sink tissues are tightly linked to leaf hydraulic traits, with little to no effect of HLB exposure duration or source:sink ratio. Some leaf hydraulic traits were correlated with fruit sucrose and fruit water content, which were primarily driven by fruit developmental stage. When fruit sucrose content was high, ψ_leaf_ was less negative and ɛ was greater, suggesting improved leaf hydraulic function. Likewise, ψ_p_ positively correlated with total fruit water loss indicating a tradeoff between leaf turgor and fruit turgor pressure, given that fruit water loss to xylem backflow is linked to reduced fruit turgor pressure (Goldschmidt & Koch 1996). This gradient promotes xylem backflow from fruit to leaves, facilitating hydraulic redistribution that may be required to support carbon export from leaves. In contrast, higher leaf *C*_tlp_ and *C\**_tlp_ and low fruit sucrose during Stage I of fruit development suggests that leaf water status maintenance may be prioritized over carbon transport during early fruit development. Consistent with these findings, Tang *et al*. (2020) reported that differences in fruit growth between severely and mildly affected ‘Valencia’ sweet oranges were evident as early as Stage I (May), and these early size differences were associated with subsequent fruit drop. Hernandez-Santana *et al*. (2021) also reported that olive fruit growth is more sensitive to leaf water status than to photosynthesis, and that this relationship varies across fruit development stages. Together, these findings suggest that xylem backflow helps maintain carbohydrate transport at the expense of fruit expansion.

## CONCLUSION

The present study reports the impacts of exposure duration of a phloem-affecting disease on leaf hydraulic properties and fruit-leaf water dynamics, revealing impacts of source-sink dysfunction on aboveground water dynamics. Prolonged exposure to HLB promotes acclimation to water deficit, modifies leaf water status, and increases leaf reliance on fruit xylem backflow. Xylem backflow is critical to maintaining phloem pressure gradient particularly under water deficit conditions. The study also highlights fruit diurnal shrinkage pattern as an efficient non-destructive and consistent tool for assessing water-carbon tradeoffs between leaves and fruits. Further studies are needed on strategies to enhance plant hydraulic function and effective carbon transport. Additionally, water management strategies should consider fruit developmental stages, as sensitivity to water deficit varies across stages.

## Author Contributions

C.V. and D.A.: conceptualization, methodology; D.A.: implementation of the study, data curation, statistical analysis, writing-original draft preparation; C.V.: supervision; C.V. and T.V.: writing-reviewing and editing. All authors have read and agreed to the published version of the manuscript.

## Supporting information

Supplementary figures supporting results

## Acknowledgements

We sincerely appreciate Dr. Lauren Diepenbrock for permission to the experimental plot and permitting us to use the site for this study. Thank you to Gillian Z. Michalczyk, Faustin Mukunzi, Myrtho Pierre, Jean Francois, Maryam Khalilzadeh, and Rashmi Kumari for all the help and support during the data collection and laboratory sample processing.

