## Supplementary figures supporting results for "PLANT WATER-CARBON RELATIONS IN HUANGLONGBING-AFFECTED CITRUS TREES AND THE IMPACT OF HLB EXPOSURE DURATION, FRUIT DEVELOPMENTAL STAGE, AND SOURCE-SINK MANIPULATION ON THESE RELATIONS"

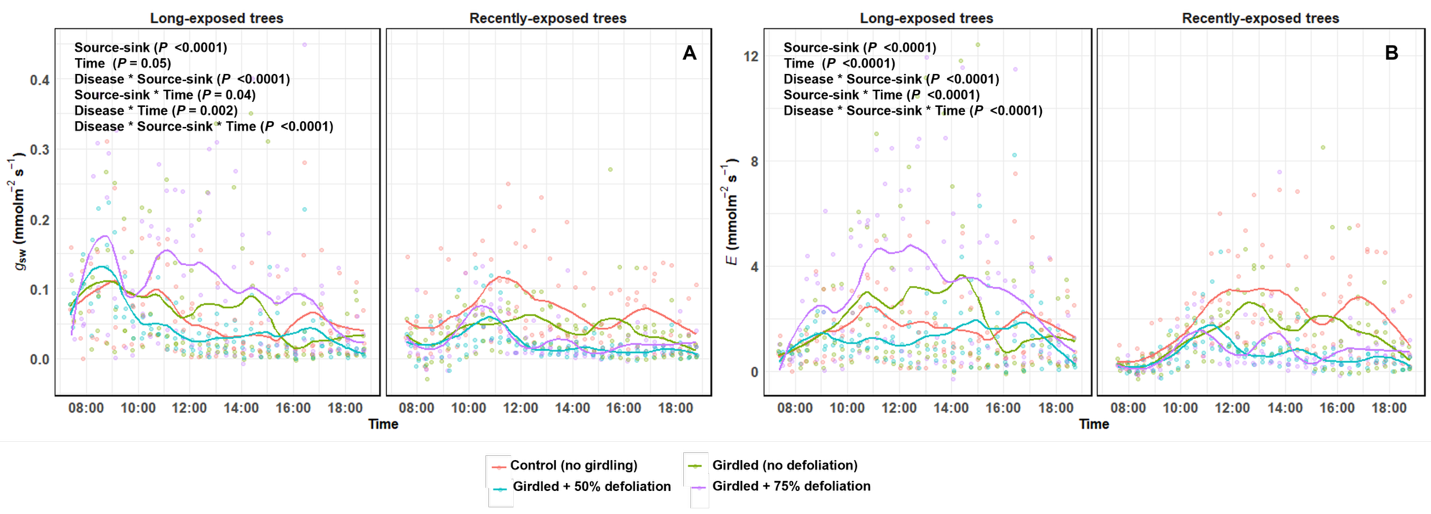

Figure S1: Diurnal leaf stomatal conductance (*g*_sw_) and transpiration (*E*) of field-grown *Citrus sinensis* 'Valencia' trees in a study of Huanglongbing (HLB) disease and various source-sink manipulations taken between 7:00 am – 7:00 pm. Data collected at sampling dates during Stage I - III of fruit development. Means labeled with different letters are significantly different as determined using Tukey’s honestly significant difference at *P* < 0.05.

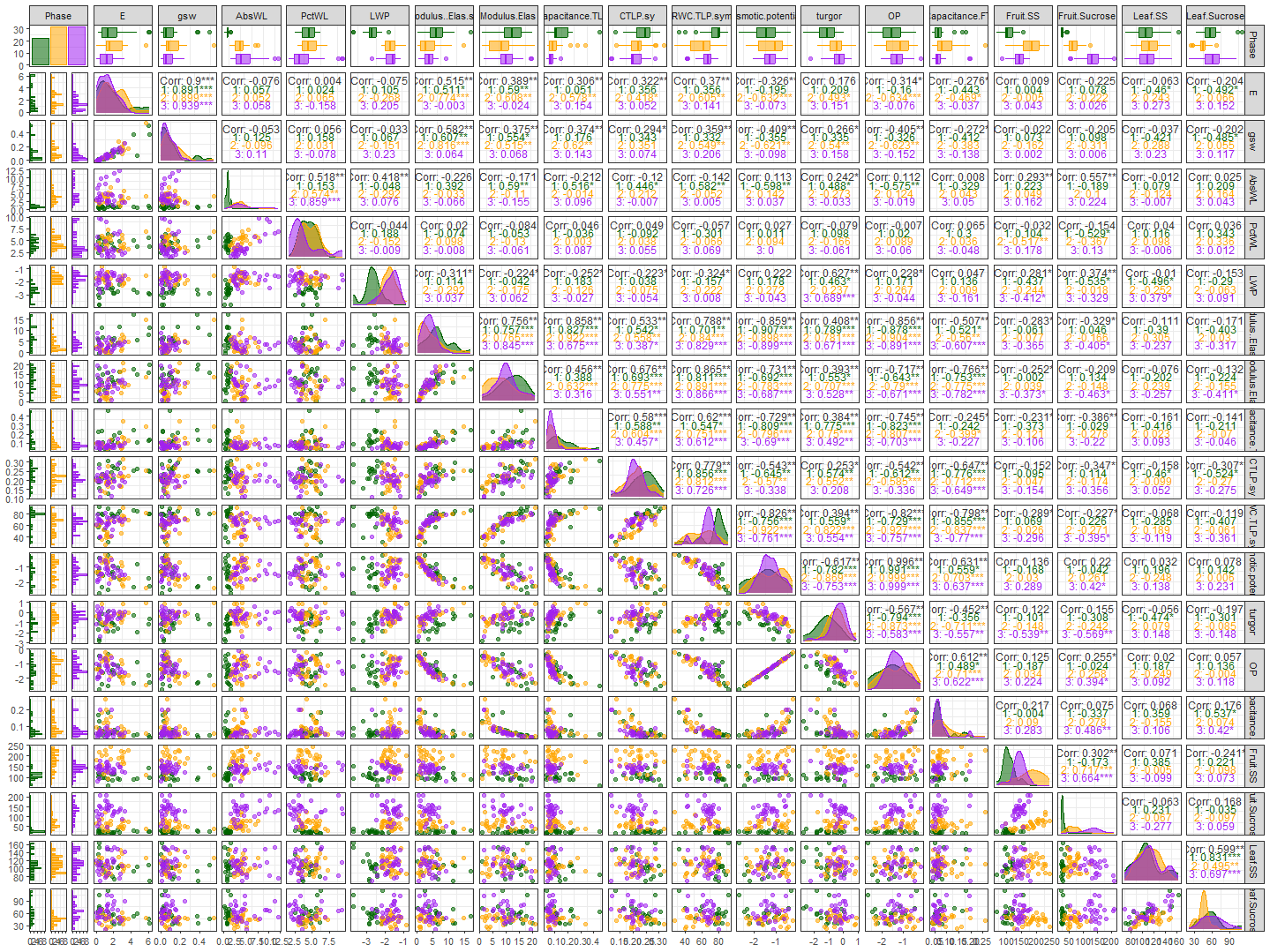

Figure S2: Leaf hydraulic traits relationship with leaf and fruit sugar content of field-grown *Citrus* x *sinensis* 'Valencia' trees in a study of Huanglongbing (HLB) disease and various source-sink manipulations during Stage I - III of fruit development. 1, 2 and 3 are Stages I, II and III of fruit development, E denotes leaf transpiration, gsw represents stomatal conductance, AbsWL represents total fruit water loss, LWP represents leaf water potential, Modulus.Elast.Sym represents symplastic modulus of elasticity, *C**_tlp_ denotes total capacitance at turgor loss point, and *C*_tlp_ denotes symplastic capacitance at turgor loss point. Fruit.SS denotes fruit soluble sugar, Leaf.SS denotes leaf soluble sugar.

Table S1: Weather patterns in Lake Alfred, FL, from May – November 2024, showing the monthly average and maximum temperature in (˚C), rainfall (cm).

| **Month** | **Average Temp (˚C)** | **Max Temp (˚C)** | **Rainfall (cm)** | **Relative Humidity (%)** |
| --- | --- | --- | --- | --- |
| **May-24** | 27.15 | 36.43 | 6.86 | 71 |
| **Jun-24** | 27.55 | 37.65 | 13.89 | 78 |
| **Jul-24** | 28.19 | 36.46 | 12.80 | 81 |
| **Aug-24** | 27.61 | 35.09 | 19.91 | 83 |
| **Sep-24** | 27.35 | 34.92 | 15.11 | 84 |
| **Oct-24** | 23.61 | 33.12 | 23.50 | 81 |
| **Nov-24** | 20.86 | 30.73 | 1.04 | 79 |

Table S2**:** Analysis of variance of leaf non-structural carbohydrates of field-grown *Citrus sinensis* 'Valencia' trees in a study of Huanglongbing (HLB) disease and various source-sink manipulations. Data collected at sampling dates during Stage I - III of fruit development. D- disease status; T- source-sink treatment; S- fruit development stages.

| **Variable** | **Effect** | **Num. DF** | **Den. DF** | **F value** | ***P* value** | **Variable** | **Num. DF** | **Den. DF** | **F value** | ***P* value** |
| --- | --- | --- | --- | --- | --- | --- | --- | --- | --- | --- |
| **Soluble sugar** | **D** | 1 | 18.0 | 29.6 | <0.001* | **Sucrose** | 1 | 18.4 | 27.7 | <0.001* |
|  | **T** | 3 | 191.6 | 0.6 | 0.5 |  | 3 | 188.6 | 0.5 | 0.6 |
|  | **S** | 2 | 191.5 | 13.5 | <0.001* |  | 2 | 189.3 | 57.5 | <0.001* |
|  | **D:T** | 3 | 191.6 | 1.8 | 0.1 |  | 3 | 188.6 | 1.2 | 0.2 |
|  | **D:S** | 2 | 191.5 | 21.0 | <0.001* |  | 2 | 189.3 | 12.9 | <0.001* |
|  | **T:S** | 6 | 191.6 | 0.6 | 0.7 |  | 6 | 188.6 | 0.3 | 0.8 |
|  | **D:S:T** | 6 | 191.6 | 0.5 | 0.7 |  | 6 | 188.6 | 0.8 | 0.5 |
| **Reducing sugar** | **D** | 1 | 18.0 | 6.6 | 0.01* |  |  |  |  |  |
|  | **T** | 3 | 188.0 | 0.8 | 0.4 |  |  |  |  |  |
|  | **S** | 2 | 188.7 | 40.5 | <0.001* |  |  |  |  |  |
|  | **D:T** | 3 | 188.0 | 0.5 | 0.6 |  |  |  |  |  |
|  | **D:S** | 2 | 188.7 | 5.6 | 0.004* |  |  |  |  |  |
|  | **T:S** | 6 | 188.0 | 0.5 | 0.7 |  |  |  |  |  |
|  | **D:S:T** | 6 | 188.0 | 0.1 | 0.9 |  |  |  |  |  |

Table S3**:** Analysis of variance of leaf water status of field-grown *Citrus sinensis* 'Valencia' trees in a study of Huanglongbing (HLB) disease and various source-sink manipulations. Data collected at sampling dates during Stage I - III of fruit development. D- disease status; T- source-sink treatment; S- fruit development stages. Turgor (or pressure) potential (Ψ_p_); Osmotic (or solute) potential (Ψ_o_); Osmotic potential at turgor loss point (π_TLP_); Osmotic potential at full turgor (π_o_); total and symplastic modulus of elasticity at full turgor (ɛ*, ɛ respectively); total and symplastic relative water content at turgor loss point (*RWC**_tlp_, *RWC*_tlp_ respectively); total and symplastic capacitance at full turgor (*C**_ft_ *, C*_ft_ respectively) and total and symplastic capacitance at turgor loss point (*C**_tlp_ *, C*_tlp_ respectively).

| **Variable** | **Effect** | **Num. DF** | **Den. DF** | **F value** | ***P* value** | **Variable** | **Num. DF** | **Den. DF** | **F value** | ***P* value** |
| --- | --- | --- | --- | --- | --- | --- | --- | --- | --- | --- |
| **SWC** | **D** | 1 | 7.9 | 0.2 | 0.6 | **ψ_leaf_** | 1 | 7.5 | 1.2 | 0.3 |
|  | **T** | 3 | 47.5 | 1.6 | 0.1 |  | 3 | 51.4 | 1.8 | 0.1 |
|  | **S** | 2 | 53.6 | 11.4 | <0.0001* |  | 2 | 57.0 | 65.1 | <0.001* |
|  | **D:T** | 3 | 47.5 | 4.4 | 0.007* |  | 3 | 51.4 | 0.3 | 0.7 |
|  | **D:S** | 2 | 53.6 | 0.1 | 0.8 |  | 2 | 57.0 | 2.9 | 0.05 |
|  | **T:S** | 6 | 47.9 | 5.9 | <0.0001* |  | 6 | 51.3 | 3.7 | 0.003* |
|  | **D:S:T** | 6 | 47.9 | 3.5 | 0.005* |  | 6 | 51.3 | 1.0 | 0.4 |
| **Ψ_o_** | **D** | 1 | 55 | 0.05 | 0.8 | **Ψ_p_** | 1 | 5.8 | 2.1 | 0.1 |
|  | **T** | 3 | 55 | 0.07 | 0.9 |  | 3 | 47.8 | 0.1 | 0.9 |
|  | **S** | 2 | 55 | 0.5 | 0.5 |  | 2 | 54.8 | 16.9 | <0.0001* |
|  | **D:T** | 3 | 55 | 1.4 | 0.2 |  | 3 | 47.8 | 1.9 | 0.1 |
|  | **D:S** | 2 | 55 | 4.3 | 0.01* |  | 2 | 54.8 | 3.3 | 0.04* |
|  | **T:S** | 6 | 55 | 2.3 | 0.04* |  | 6 | 48.0 | 3.6 | 0.004* |
|  | **D:S:T** | 6 | 55 | 2.2 | 0.05 |  | 6 | 48.0 | 1.0 | 0.3 |
| **π_o_** | **D** | 1 | 54 | 0.02 | 0.8 | **π_TLP_** | 1 | 54 | 0.3 | 0.5 |
|  | **T** | 3 | 54 | 0.2 | 0.8 |  | 3 | 54 | 0.08 | 0.9 |
|  | **S** | 2 | 54 | 0.2 | 0.8 |  | 2 | 54 | 0.2 | 0.8 |
|  | **D:T** | 3 | 54 | 1.8 | 0.1 |  | 3 | 54 | 1.5 | 0.2 |
|  | **D:S** | 2 | 54 | 6.0 | 0.004* |  | 2 | 54 | 5.4 | 0.006* |
|  | **T:S** | 6 | 54 | 2.7 | 0.02* |  | 6 | 54 | 1.8 | 0.1 |
|  | **D:S:T** | 6 | 54 | 2.4 | 0.03* |  | 6 | 54 | 1.9 | 0.08 |
| **a_f_ (%)** | **D** | 1 | 52 | 0.2 | 0.5 | ***RWC**_tlp_** | 1 | 4.9 | 9.5 | 0.02* |
|  | **T** | 3 | 52 | 0.8 | 0.4 |  | 3 | 45.7 | 1.1 | 0.3 |
|  | **S** | 2 | 52 | 1.8 | 0.1 |  | 2 | 53.8 | 2.9 | 0.06 |
|  | **D:T** | 3 | 52 | 0.06 | 0.9 |  | 3 | 45.7 | 0.5 | 0.6 |
|  | **D:S** | 2 | 52 | 3.7 | 0.03* |  | 2 | 53.8 | 2.8 | 0.06 |
|  | **T:S** | 6 | 52 | 0.2 | 0.9 |  | 6 | 46.3 | 1.3 | 0.2 |
|  | **D:S:T** | 6 | 52 | 1.2 | 0.2 |  | 6 | 46.3 | 1.0 | 0.4 |
| **ɛ*** | **D** | 1 | 54 | 3.7 | 0.05 | ***C**_ft_** | 1 | 54 | 3.6 | 0.06 |
|  | **T** | 3 | 54 | 1.2 | 0.3 |  | 3 | 54 | 1.1 | 0.3 |
|  | **S** | 2 | 54 | 1.2 | 0.3 |  | 2 | 54 | 2.2 | 0.1 |
|  | **D:T** | 3 | 54 | 1.2 | 0.3 |  | 3 | 54 | 0.8 | 0.4 |
|  | **D:S** | 2 | 54 | 5.8 | 0.004* |  | 2 | 54 | 6.0 | 0.004* |
|  | **T:S** | 6 | 54 | 1.6 | 0.1 |  | 6 | 54 | 2.3 | 0.04* |
|  | **D:S:T** | 6 | 54 | 0.7 | 0.6 |  | 6 | 54 | 0.8 | 0.5 |
| ***C**_tlp_** | **D** | 1 | 10.0 | 0.09 | 0.9 | **ɛ** | 1 | 50 | 0.1 | 0.7 |
|  | **T** | 3 | 48.6 | 0.09 | 0.9 |  | 3 | 50 | 0.5 | 0.6 |
|  | **S** | 2 | 52.9 | 4.6 | 0.01* |  | 2 | 50 | 2.1 | 0.1 |
|  | **D:T** | 3 | 48.6 | 0.5 | 0.6 |  | 3 | 50 | 1.2 | 0.3 |
|  | **D:S** | 2 | 52.9 | 1.1 | 0.3 |  | 2 | 50 | 6.3 | 0.03* |
|  | **T:S** | 6 | 49.3 | 1.2 | 0.3 |  | 6 | 50 | 2.9 | 0.01* |
|  | **D:S:T** | 6 | 49.3 | 1.1 | 0.3 |  | 6 | 50 | 1.5 | 0.1 |
| ***RWC*_tlp_** | **D** | 1 | 54 | 2.8 | 0.09 | ***C*_tlp_** | 1 | 8.1 | 4.4 | 0.06 |
|  | **T** | 3 | 54 | 1.8 | 0.1 |  | 3 | 48.0 | 2.1 | 0.1 |
|  | **S** | 2 | 54 | 2.1 | 0.1 |  | 2 | 53.8 | 2.0 | 0.1 |
|  | **D:T** | 3 | 54 | 1.0 | 0.3 |  | 3 | 48.0 | 0.3 | 0.8 |
|  | **D:S** | 2 | 54 | 9.3 | 0.0003* |  | 2 | 53.8 | 3.17 | 0.04* |
|  | **T:S** | 6 | 54 | 3.2 | 0.008* |  | 6 | 48.4 | 1.5 | 0.1 |
|  | **D:S:T** | 6 | 54 | 1.6 | 0.2 |  | 6 | 48.4 | 0.8 | 0.5 |
| ***C*_ft_** | **D** | 1 | 54 | 0.6 | 0.4 |  |  |  |  |  |
|  | **T** | 3 | 54 | 1.5 | 0.2 |  |  |  |  |  |
|  | **S** | 2 | 54 | 0.7 | 0.4 |  |  |  |  |  |
|  | **D:T** | 3 | 54 | 1.3 | 0.2 |  |  |  |  |  |
|  | **D:S** | 2 | 54 | 8.2 | <0.001* |  |  |  |  |  |
|  | **T:S** | 6 | 54 | 3.1 | 0.01* |  |  |  |  |  |
|  | **D:S:T** | 6 | 54 | 1.5 | 0.1 |  |  |  |  |  |

Table S4**:** Analysis of variance of midday and diurnal stomatal conductance (*g*_sw_) and transpiration (*E*) of field-grown *Citrus sinensis* 'Valencia' trees in a study of Huanglongbing (HLB) disease and various source-sink manipulations. Data collected at sampling dates during Stage I - III of fruit development. D represents disease status; T represents source-sink treatment; S represents fruit development stages.

| **Variable** | **Effect** | **Num. DF** | **Den. DF** | **F value** | ***P* value** | **Variable** | **Num. DF** | **Den. DF** | **F value** | ***P* value** |
| --- | --- | --- | --- | --- | --- | --- | --- | --- | --- | --- |
| **Midday *g*_sw_** | **T** | 3 | 820 | 11.6 | <0.0001* | **Midday *E*** | 3 | 820 | 13.9 | <0.0001* |
|  | **D** | 1 | 4 | 3.1 | 0.1 |  | 1 | 4 | 1.9 | 0.2 |
|  | **Ti** | 1 | 820 | 3.6 | 0.05 |  | 1 | 820 | 74.9 | <0.0001* |
|  | **D:T** | 3 | 820 | 17.2 | <0.0001* |  | 3 | 820 | 21.6 | <0.0001* |
|  | **T:Ti** | 3 | 820 | 2.6 | 0.04* |  | 3 | 820 | 3.4 | 0.01* |
|  | **D:Ti** | 1 | 820 | 9.5 | 0.002* |  | 1 | 820 | 0.3 | 0.5 |
|  | **D:Ti:T** | 3 | 820 | 3.3 | 0.01* |  | 3 | 820 | 5.6 | <0.0001* |
| **Diurnal *g*_sw_** | **D** | 1 | 18 | 6.3 | 0.02* | **Diurnal *E*** | 1 | 18 | 7.8 | 0.01* |
|  | **T** | 3 | 198 | 2.2 | 0.07 |  | 3 | 198 | 2.8 | 0.03* |
|  | **S** | 2 | 198 | 4.0 | 0.01* |  | 2 | 198 | 5.3 | 0.005* |
|  | **D:T** | 3 | 198 | 0.2 | 0.8 |  | 3 | 198 | 0.7 | 0.5 |
|  | **D:S** | 2 | 198 | 11.0 | <0.0001* |  | 2 | 198 | 13.1 | <0.0001* |
|  | **T:S** | 6 | 198 | 1.0 | 0.3 |  | 6 | 198 | 0.5 | 0.8 |
|  | **D:S:T** | 6 | 198 | 1.0 | 0.4 |  | 6 | 198 | 0.6 | 0.6 |

Table S5**:** Analysis of variance of fruit non-structural carbohydrates of field-grown *Citrus sinensis* 'Valencia' trees in a study of Huanglongbing (HLB) disease and various source-sink manipulations. Data collected at sampling dates during Stage I - III of fruit development. D represents disease status; T represents source-sink treatment; S represents fruit development stages.

| **Variable** | **Effect** | **Num. DF** | **Den. DF** | **F value** | ***P* value** | **Variable** | **Num. DF** | **Den. DF** | **F value** | ***P* value** |
| --- | --- | --- | --- | --- | --- | --- | --- | --- | --- | --- |
| **Soluble sugar** | **D** | 1 | 18.0 | 0.3 | 0.5 | **Sucrose** | 1 | 17.2 | 1.3 | 0.2 |
|  | **T** | 3 | 192.4 | 0.5 | 0.6 |  | 3 | 192.0 | 1.0 | 0.3 |
|  | **S** | 2 | 192.4 | 137.1 | <0.001* |  | 2 | 192.0 | 488.1 | <0.001* |
|  | **D:T** | 3 | 192.4 | 0.6 | 0.6 |  | 3 | 192.0 | 1.0 | 0.3 |
|  | **D:S** | 2 | 192.4 | 11.1 | <0.001* |  | 2 | 192.0 | 3.3 | 0.03* |
|  | **T:S** | 6 | 192.4 | 0.4 | 0.8 |  | 6 | 192.0 | 0.5 | 0.7 |
|  | **D:S:T** | 6 | 192.4 | 0.6 | 0.6 |  | 6 | 192.0 | 0.4 | 0.8 |
| **Reducing sugar** | **D** | 1 | 18.1 | 0.02 | 0.8 |  |  |  |  |  |
|  | **T** | 3 | 192.5 | 0.4 | 0.7 |  |  |  |  |  |
|  | **S** | 2 | 192.5 | 355.2 | <0.001* |  |  |  |  |  |
|  | **D:T** | 3 | 192.5 | 1.5 | 0.2 |  |  |  |  |  |
|  | **D:S** | 2 | 192.5 | 3.9 | 0.02* |  |  |  |  |  |
|  | **T:S** | 6 | 192.5 | 0.5 | 0.7 |  |  |  |  |  |
|  | **D:S:T** | 6 | 192.5 | 0.4 | 0.8 |  |  |  |  |  |

Table S6**:** Analysis of variance for diurnal fruit shrinkage of field-grown *Citrus sinensis* 'Valencia' trees in a study of Huanglongbing (HLB) disease and various source-sink manipulations. Data collected at sampling dates during Stage I - III of fruit development. D represents disease status; T represents source-sink treatment; S represents fruit development stages; Ti represents time of day.

| **Stages** | **Effect** | **Num. DF** | **Den. DF** | **F value** | ***P* value** | **Stages** | **Num. DF** | **Den. DF** | **F value** | ***P* value** |
| --- | --- | --- | --- | --- | --- | --- | --- | --- | --- | --- |
| **Stage 1** | **D** | 1 | 80.26 | 7.04 | 0.009* | **Stage 2** | 1 | 18 | 4.1 | 0.05 |
|  | **T** | 3 | 69.09 | 0.9 | 0.4 |  | 3 | 54 | 0.3 | 0.7 |
|  | **Ti** | 4 | 2197.97 | 412.2 | <0.001* |  | 4 | 2288 | 612.5 | <0.001* |
|  | **D:T** | 3 | 82.13 | 1.18 | 0.3 |  | 3 | 54 | 0.3 | 0.7 |
|  | **D:Ti** | 4 | 2196.15 | 14.5 | <0.001* |  | 4 | 2288 | 21.8 | <0.001* |
|  | **T:Ti** | 12 | 2197.5 | 11.8 | <0.001* |  | 12 | 2288 | 6.15 | <0.001* |
|  | **D:Ti:T** | 12 | 2196.0 | 5.1 | <0.001* |  | 12 | 2288 | 3.9 | <0.001* |
| **Stage 3** | **D** | 1 | 17.7 | 0.8 | 0.3 |  |  |  |  |  |
|  | **T** | 3 | 54.2 | 0.06 | 0.9 |  |  |  |  |  |
|  | **Ti** | 4 | 2258.2 | 218.0 | <0.001* |  |  |  |  |  |
|  | **D:T** | 3 | 54.2 | 0.1 | 0.9 |  |  |  |  |  |
|  | **D:Ti** | 4 | 2258.2 | 25.0 | <0.001* |  |  |  |  |  |
|  | **T:Ti** | 12 | 2258.2 | 5.4 | <0.001* |  |  |  |  |  |
|  | **D:Ti:T** | 12 | 2258.2 | 7.5 | <0.001* |  |  |  |  |  |

Table S7**:** Analysis of variance for absolute and relative fruit water loss of field-grown *Citrus sinensis* 'Valencia' trees in a study of Huanglongbing (HLB) disease and various source-sink manipulations. Data collected at sampling dates during Stage I - III of fruit development. D represents disease status; T represents source-sink treatment; Abs denotes absolute fruit water loss; Pct denotes relative fruit water loss.

| **Stages** | **Variable** | **Effect** | **Num. DF** | **Den. DF** | **F value** | ***P* value** | **Variable** | **Num. DF** | **Den. DF** | **F value** | ***P* value** |
| --- | --- | --- | --- | --- | --- | --- | --- | --- | --- | --- | --- |
| **1** | **Abs. H_2_O loss** | **D** | 1 | 25.6 | 5.6 | 0.02* | **Pct H_2_O loss** | 1 | 28.2 | 0.5 | 0.4 |
|  |  | **T** | 3 | 55.9 | 0.5 | 0.6 |  | 3 | 56.0 | 1.1 | 0.3 |
|  |  | **D:T** | 3 | 56.0 | 0.4 | 0.7 |  | 3 | 56.2 | 0.7 | 0.5 |
| **2** | **Abs. H_2_O loss** | **D** | 1 | 72 | 0.3 | 0.5 | **Pct H_2_O loss** | 1 | 72 | 6.1 | 0.01* |
|  |  | **T** | 3 | 72 | 0.09 | 0.9 |  | 3 | 72 | 0.2 | 0.8 |
|  |  | **D:T** | 3 | 72 | 1.2 | 0.2 |  | 3 | 72 | 0.4 | 0.7 |
| **3** | **Abs. H_2_O loss** | **D** | 1 | 17.9 | 0.0 | 0.9 | **Pct H_2_O loss** | 1 | 17.9 | 7.6 | 0.02* |
|  |  | **T** | 3 | 54.4 | 0.3 | 0.7 |  | 3 | 54.5 | 0.03 | 0.9 |
|  |  | **D:T** | 3 | 54.4 | 1.1 | 0.3 |  | 3 | 54.5 | 0.4 | 0.7 |

Table S8**:** Analysis of variance for diurnal fruit water loss relative to fruit water content of field-grown *Citrus sinensis* 'Valencia' trees in a study of Huanglongbing (HLB) disease and various source-sink manipulations. Data collected at sampling dates during Stage I - III of fruit development. D represents disease status; T represents source-sink treatment; S represents fruit development stages; Ti represents time of day.

| **Effect** | **Num. DF** | **Den. DF** | **F value** | ***P* value** |
| --- | --- | --- | --- | --- |
| **D** | 1 | 18.9 | 3.4 | 0.07 |
| **T** | 3 | 117.0 | 2.1 | 0.1 |
| **Ti** | 1 | 123.7 | 1.1 | 0.2 |
| **D:T** | 3 | 117.0 | 0.3 | 0.7 |
| **D:Ti** | 1 | 123.7 | 0.1 | 0.6 |
| **T:Ti** | 3 | 116.9 | 0.05 | 0.9 |
| **D:Ti:T** | 3 | 116.9 | 1.3 | 0.2 |

Table S9: Linear regression coefficients of field-grown *Citrus sinensis* 'Valencia' trees in a study of Huanglongbing (HLB) disease and various source-sink manipulations. Data collected at sampling dates during Stage I - III of fruit development. *C*_tlp_ and *C**_tlp_ represents symplastic and total capacitance at turgor loss point respectively; ψ_leaf_ represents leaf water potential; ɛ denotes symplastic modulus of elasticity.

| **Variable** | **Effect** | **Df** | **F value** | ***P* value** | ***R*^2^** |
| --- | --- | --- | --- | --- | --- |
| **Leaf sucrose** | ***C*_tlp_** | 1 | 8.8 | 0.004* | 0.32 |
|  | **Stage** | 2 | 10.1 | 0.0001* |  |
|  | ***C*_tlp_:Stage** | 2 | 2.0 | 0.1 |  |
| **Fruit sucrose** | **ψ_leaf_** | 1 | 37.0 | <0.0001* | 0.73 |
|  | **Stage** | 2 | 83.7 | <0.0001* |  |
|  | **ψ_leaf_:Stage** | 2 | 1.5 | 0.2 |  |
| **Fruit sucrose** | **ɛ** | 1 | 21.4 | <0.0001* | 0.75 |
|  | **Stage** | 2 | 90.1 | <0.0001* |  |
|  | **ɛ:Stage** | 2 | 3.7 | 0.02* |  |
| **Fruit sucrose** | ***C**_tlp_** | 1 | 33.6 | <0.0001* | 0.73 |
|  | **Stage** | 2 | 79.0 | <0.0001* |  |
|  | ***C**_tlp_:Stage** | 2 | 1.0 | 0.3 |  |
| **Fruit sucrose** | ***C*_tlp_** | 1 | 31.3 | <0.0001* | 0.75 |
|  | **Stage** | 2 | 90.7 | <0.0001* |  |
|  | ***C*_tlp_:Stage** | 2 | 2.2 | 0.1 |  |
